# Dopamine facilitates the response to glutamatergic inputs in a computational model of astrocytes

**DOI:** 10.1101/2022.11.10.516040

**Authors:** Thiago Ohno Bezerra, Antonio C. Roque

## Abstract

Astrocytes are active cells that respond to neurotransmitters with elevations in their intracellular calcium concentration (calcium signals). In a tripartite synapse involving two neurons coupled by a glutamatergic synapse and one astrocyte, glutamate released by the presynaptic neuron can generate calcium signals in the astrocyte, which in turn trigger the release of neuroactive molecules (gliotransmitters) by the astrocyte that bind to receptors in the pre- and postsynaptic neuron membranes and modulate synaptic transmission. Astrocytic calcium signals can also be evoked by dopamine released in distant sites. Little is known about how dopamine modulates glutamatergic-evoked astrocyte activity. To investigate this question, we constructed compartmental astrocyte models with three different morphologies: linear (soma plus a single branch); branched (soma plus two branches); and bifurcated (soma plus a single branch that bifurcates into two branchlets). Compartments were modeled by conductance-based equations for membrane voltage and transport of ions, glutamate and dopamine between extra- and intracellular spaces. Glutamatergic and dopaminergic stimuli were modeled as Poisson processes with variable frequencies, and astrocyte responses were measured by number and location of evoked calcium signals. For cells with linear morphology, whole-cell dopaminergic stimulation reduced the glutamatergic stimulation frequency of distal compartments needed to generate calcium signals. For both the branched and bifurcated morphologies, whole-cell dopaminergic stimulation together with glutamatergic stimulation of one of the processes reduced the glutamatergic stimulation frequency necessary to trigger a calcium signal in the other process. The same glutamatergic stimulation protocols without dopamine stimulation required higher glutamatergic input frequencies to evoke calcium signals. Our results suggest that dopamine facilitates the occurrence of glutamatergic-evoked calcium signals, and that dopamine-glutamate interaction can control the distribution of calcium signals along the astrocyte extension.

**Author summary:** Astrocytes are brain cells that are not electrically excitable as neurons but display chemical excitability in the form of transient rises in the intracellular calcium concentration (calcium signals) evoked by neurotransmitters. A tripartite synapse consists of pre- and postsynaptic terminals ensheathed by astrocyte processes. Neurotransmitters released by the presynaptic neuron can generate calcium signals in the astrocyte, which in turn trigger the release of neuroactive molecules (gliotransmitters) by the astrocyte that bind to receptors in the pre- and postsynaptic membranes and modulate synaptic transmission. Two neurotransmitters that can evoke astrocytic calcium signals are glutamate, the major neurotransmitter of excitatory synapses, and dopamine, an important modulatory neurotransmitter that can diffuse to wider regions than the synaptic release site. Little is known about how dopamine modulates glutamatergic-evoked astrocyte activity, and here we investigate this question using computational modeling. We constructed compartmental astrocyte models with three different morphologies: linear, with a single branch emanating from soma; branched, with two branches emanating from soma; and bifurcated, with a branch emanating from soma that bifurcates into two branchlets. Compartments were modeled by conductance-based equations for membrane voltage and transport of ions (sodium, potassium and calcium), glutamate and dopamine between extra- and intracellular spaces. Glutamatergic and dopaminergic stimuli were modeled as Poisson processes with variable frequencies. Astrocyte models with the three morphologies were submitted to similar stimulation protocols to compare their responses, which were measured in terms of the frequency and location of evoked calcium signals. For cells with linear morphology, dopaminergic stimulation of the entire cell (to simulate the diffuse action of dopamine) reduced the glutamatergic stimulation frequency of distal compartments (which simulates glutamatergic input from presynaptic neuron) needed to generate calcium signals. For both the branched and bifurcated morphologies, dopaminergic stimulation of the whole cell together with glutamatergic stimulation of the distal portions of one of the processes reduced the glutamate stimulation frequency necessary to trigger a calcium signal in the distal portions of the other process. Repetitions of the glutamatergic stimulation protocols without whole cell dopaminergic stimulation showed that higher glutamatergic input frequencies were needed to evoke calcium signals. Our results suggest that dopamine facilitates the occurrence of calcium signals evoked by glutamatergic inputs, and that interaction between dopamine and glutamate can control the distribution of calcium signals along the astrocyte extension.

## Introduction

Classically, astrocytes were associated with neuronal support functions like potassium buffering, metabolic supply and neurotransmitter reuptake [1]. However, apart from their classical role, recent studies have shown that astrocytes are related to brain and cognitive functions like working memory, long-term memory and goal-directed behavior [2–4]. Some studies have even suggested that the behavioral impairments caused by cannabidiol and amphetamine administration in mice are dependent on astrocytic activity [2, 7].

Astrocytes are active cells that respond to neurotransmitters with an increase in the intracellular Ca^2+^ concentration [5, 6, 8], a ‘calcium signal’. The elevation in the intracellular Ca^2+^ concentration promotes the release of neuroactive molecules called gliotransmitters, e.g. glutamate, ATP and D-serine [1]. Release of gliotransmitters modulate synaptic transmission, increase neuron excitability, influence synaptic plasticity, and enhance neural synchronization [3, 5]. Fine astrocytic processes ensheath synaptic terminals, forming a so-called tripartite synapse. Glutamate released by the presynaptic cell in a tripartite synapse is the major source of glutamatergic input to the astrocyte [19]. However, neuromodulators such as dopamine and noradrenaline also promote astrocytic activation [7, 11]. Since neuromodulators act by volume transmission due to diffusion in the extracellular medium, they are particularly suited to modulate astrocytic activity [18, 20]. On the other hand, glutamatergic input could be associated to local activity in astrocytic branches. As shown in Bindocci *et al*. [17], astrocytes can show two spatially distinct activation patterns: a locally restricted activity and a global increase in the intracellular calcium associated with independent activation of several astrocytic branches. Contradicting previous assumptions, Bindocci and colleagues showed that astrocytic activity is rather local, with Ca^2+^ signals evoked in distal parts rarely reaching the soma. Global events are probably linked to stimuli reaching extensive astrocytic regions. A question that remains open is how the global events are triggered and what is their function.

Glutamate can evoke a Ca^2+^ signal by two independent mechanisms [8, 10]. Activation of glutamatergic metabotropic receptors (mGluR) expressed on the astrocytic membrane induce the synthesis of IP_3_ by phospholipase C (PLC) [8, 9]. IP_3_ activates IP_3_ receptors on the endoplasmic reticulum (ER) membrane, releasing Ca^2+^ from ER internal stores and increasing cytosolic Ca^2+^ concentration. In order to maintain the ER Ca^2+^ stores, intracellular Ca^2+^ is pumped back into the ER by the sarco/endoplasmic reticulum Ca^2+^-ATPase (SERCA) [1]. Ca^2+^ leakage through channels present in the ER membrane also influence the astrocytic Ca^2+^ concentration. A second mechanism whereby glutamate can influence the astrocytic Ca^2+^ concentration is by activating the glutamate transporter (GluT) in conjunction with the Na^+^/Ca^2+^-exchanger (NCX) [8, 10]. Na^+^ is transported into the astrocyte in each GluT cycle. To maintain Na^+^ stable, NCX promotes eflux of Na^+^ and influx of Ca^2+^.

The above two glutamatergic mechanisms of Ca^2+^ signal generation show spatial separation [10]. While the mGluR-dependent mechanism is dominant in astrocytic regions with high ER-cytosol volume ratio [16], the GluT-dependent mechanism is stronger in fine astrocytic processes that have low ER volume compared to the soma or proximal regions in astrocytic processes. Since the main glutamatergic input to an astrocyte is linked to tripartite synapses, the GluT-dependent mechanism can play a major role on glutamatergic-evoked Ca^2+^ signals.

Similarly, dopamine was shown to increase the intracellular Ca^2+^ concentration by activating D_1_ receptors expressed on the astrocytic membrane. As mGluR, these receptors activate the IP_3_-PLC pathway [7, 11], leading to Ca^2+^ efflux from ER stores. Since astrocytic fine processes show lower ER volume and no dopaminergic mechanism for Ca^2+^ generation, it is likely that dopaminergic transmission plays a modulatory role over the astrocytic activity. However, it is not yet clear what function dopamine plays in modulating the activity of astrocytes.

Although these questions are important to characterize the astrocyte function and its role in neural systems, it is difficult to monitor a whole astrocyte and stimulate astrocytic processes in specific regions. This calls for the use of computational and mathematical modeling techniques to help elucidating these questions. They offer the benefit to study and control the stimulation of specific astrocytic portions while monitoring the responses in different regions.

Li and Rinzel [13] developed a two variable model that describes the relationship between intracellular Ca^2+^ and IP_3_. Based on this model, De Pitt’a and colleagues [9] implemented a model for the mGluR-dependent mechanism of glutamatergic influence over astrocytic intracellular Ca^2+^. In that model, mGluR activates the IP_3_ pathway and lead to the efflux of Ca^2+^ from the ER. They showed that astrocytes can encode information by frequency or amplitude modes, depending on the values of the model parameters.

More recently, Oschmann and colleagues [10] included the GluT mechanism of glutamatergic influence over intracellular Ca^2+^ in their astrocyte model. They also described how these two mechanisms are related to the ER-cytosol volume ratio, showing that in fine processes the GluT-mechanism dominates the intracellular Ca^2+^ dynamics, while the mGluR mechanism is the main driving factor evoking Ca^2+^ signals in thicker branches. Other works studied the astrocyte function by using compartmental models [14] or detailed data-driven spatial templates [24]. In particular, Gordleeva *et al*. [14] showed that gliotransmitters released by astrocytes can potentiate synaptic activity and synchronize networks of neurons.

However, to the best of our knowledge, no model has yet investigated how dopaminergic input can modulate astrocytic response to glutamatergic stimulation. Since dopamine can modulate neural activity and impact astrocytic activity [11, 25], understanding its joint effect with glutamate on astrocytic activity is essential to unravel its role in regulating brain function. In this work we investigate using astrocyte compartmental models with three different morphologies (two of them branched) how glutamate and dopamine can interact and affect the intracellular Ca^2+^ concentration. We study how these neurotransmitters can be related to global events described by Bindocci *et al*. [17] and what could be the function of these global events. Initially, we show that both glutamate and dopamine can generate Ca^2+^ signals by an IP_3_-dependent mechanism and, in the case of glutamate, also by a GluT-dependent mechanism. Then we show that both the glutamatergic stimulation of the somatic compartment or the dopaminergic stimulation of the entire astrocyte enhance the response to glutamatergic inputs arriving at distal compartments, which represent glutamatergic synapses. For the branched morphologies, we also show that the joint effect of glutamate and dopamine can enable the communication between different astrocytic branches.

## Methods

### Model Description

The following variables were considered in the astrocyte model: intracellular and extracellular Ca^2+^ concentrations ([Ca^2+^]_*i*_ and [Ca^2+^]_*e*_, respectively); ER Ca^2+^ concentration ([Ca^2+^]_ER_); intracellular IP_3_ concentration ([IP_3_]); intracellular and extracellular Na^+^ concentrations ([Na^+^]_*i*_ and [Na^+^]_*e*_, respectively); intracellular and extracellular K^+^ concentrations ([K^+^]_*i*_ and [K^+^]_*e*_, respectively); the compartment potential (*V*); and extracellular glutamate ([Glu]) and dopamine ([DA]) concentrations. The equations governing each variable in the model are described in the Section “Astrocyte Model Equations”.

The compartmental model of the astrocyte is composed of a spherical soma with interconnected unit-length cylindrical compartments representing the astrocytic processes (Fig 1). The morphological parameters of the astrocyte model are based on experimental data [15]. Three different morphologies were considered in the present work, as described in the Section “Astrocyte Morphology”.

**Fig 1.**
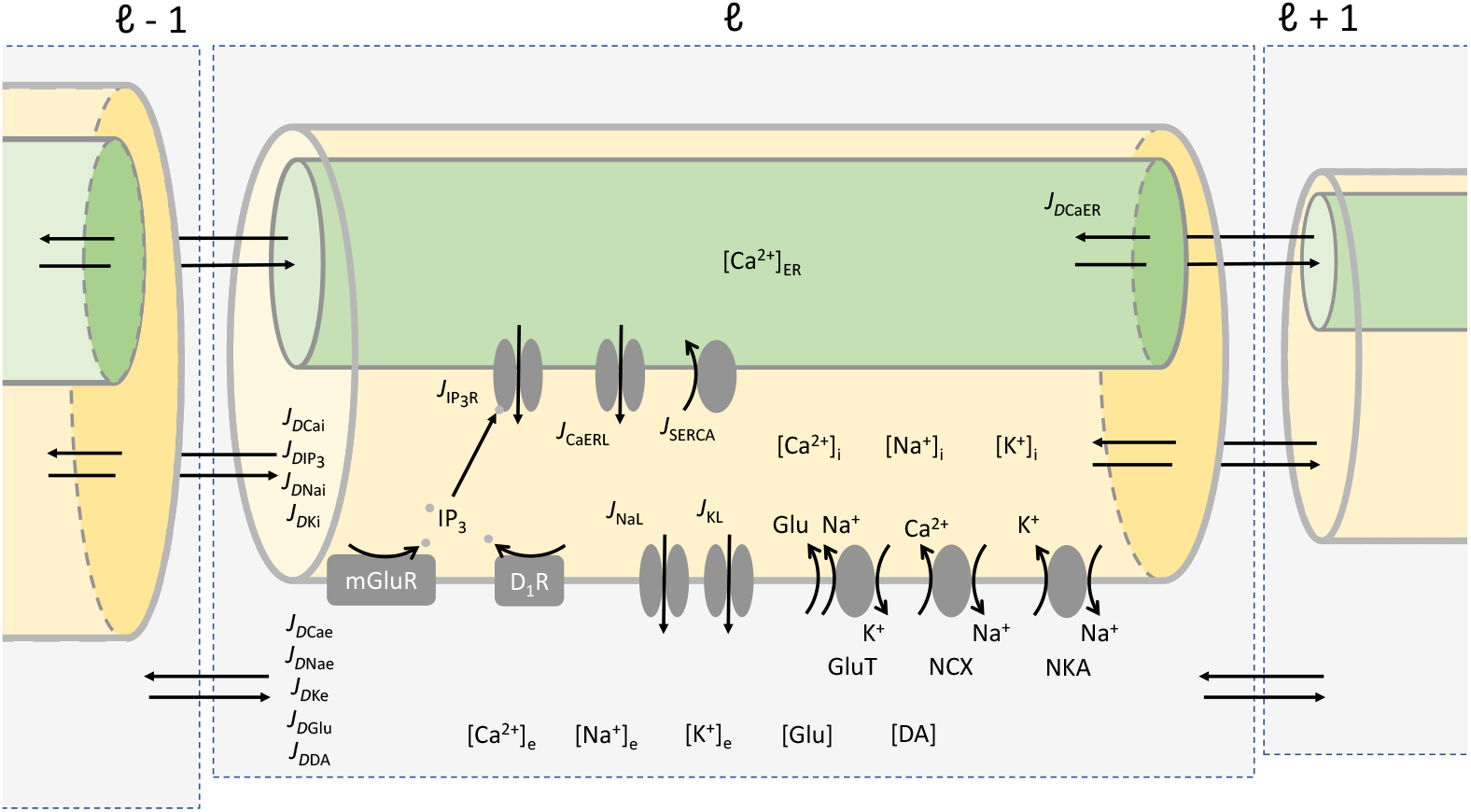
Generic astrocyte compartment showing the model variables. Each compartment is subdivided into intracellular space (yellow cylinder), ER (green cylinder) and extracellular space (gray box). The intracellular Ca^2+^ concentration is influenced by activation of IP_3_ receptors in the ER membrane, Ca^2+^ leakage from ER, SERCA pump uptake into ER, current through NCX and Ca^2+^ diffusion between intracellular compartments (*J*_*D*Cai_). The Ca^2+^ concentration in ER is governed by activation of IP_3_ receptors, leakage from ER, SERCA pump into ER and Ca^2+^ diffusion between ER compartments (*J*_*D*CaER_). The extracellular Ca^2+^ concentration is affected by the current through NCX and Ca^2+^ diffusion between extracellular compartments (*J*_*D*Cae_). The IP_3_ concentration is determined by the activation of mGluR and D_1_ receptors and the IP_3_ diffusion between intracellular compartments 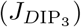. Extracellular glutamate ([Glu]) also promotes influx of Na^+^ and efflux of K^+^ through a GluT-dependent mechanism. The intracellular and extracellular Na^+^ concentrations are determined by the leak current density through Na^+^ channels (*J*_NaL_), the pump current through NKA and by Na^+^ diffusion (intracellular: *J*_*D*Nai_; extracellular: *J*_*D*Nae_). Similarly, the intracellular and extracellular K^+^ concentrations are controlled by the K^+^ leak current density (*J*_KL_), the NKA current and K^+^ diffusion (intracellular: *J*_*D*Ki_; extracellular: *J*_*D*Ke_). The extracellular concentrations of glutamate ([Glu]) and dopamine ([DA]) are affected by glutamate and dopamine diffusion in the extracellular space (*J*_*D*Glu_ and *J*_*D*DA_, respectively).

Each compartment is subdivided into three parts: a) intracellular space; b) extracellularspace space; and c) ER. These parts are interconnected by diffusion currents (Fig 1). The intra- and extracellular spaces have equal volumes. Since the volume of the ER varies along the astrocytic processes, in order to adjust the current that flows through the ER to the intracellular space, we implemented a factor *r*_ER_ representing the ratio between the ER and the cytosol volumes [10, 14]. This ratio is dependent on the surface-to-volume ratio of each compartment, *A/*𝒱, where *A* and 𝒱 are the area and volume of the compartment, respectively [16]. The factor *r*_ER_ is calculated as:

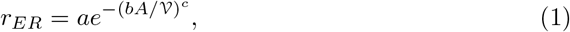

where the parameters *a* = 0.15, *b* = 0.073 μm and *c* = 2.34 were fitted using experimental data [16].

There are two distinct spatial glutamatergic mechanisms that can influence the intracellular Ca^2+^ concentration in astrocytes [10]. These mechanisms were also implemented in the present compartmental model. In proximal compartments, since the value of *r*_ER_ is higher, the mechanism that dominates the Ca^2+^ signal generation involves the activation of mGluR and the IP_3_ pathway. On the other hand, in distal compartments, which have lower *r*_ER_ values, the dominant glutamatergic mechanism involves the currents through the glutamatergic transporter (GluT), the Na^+^/Ca^2+^-exchanger (NCX) and the Na^+^/K^+^-pump (NKA) [10]. Similarly, dopamine activates the D_1_ receptors on the astrocyte membrane and the IP_3_ pathway [11]. These mechanism are described below (Eqs. 11, 22 and 12).

### Astrocyte Model Equations

For convenience, the units of the current densities *J*_NCX_, *J*_NKA_, *J*_GluT_, 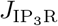, *J*_SERCA_, *J*_CaERL_, *J*_NaL_, *J*_KL_ were chosen as pA/μm^2^. This was done to simplify the calculation of the compartment potential *V* (Eq 9). In order to calculate the rates of change of molar concentrations (in μM/s), current densities were multiplied by the factor (*A/*𝒱*F*), where *F* is the Faraday constant (*F* = 96,500 C/mol). The equations below refer to a given compartment ℓ. However, in order to not overload the equations we omitted the compartment index. All diffusion current densities are modeled as indicated in Eq (27).

The intracellular Ca^2+^ concentration is given by [9, 14]:

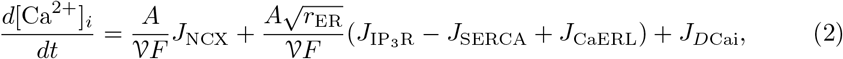

where *J*_NCX_ is the NCX current density (Eq 24), 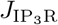 is the current density through the IP_3_R channel in the ER (Eq 18), *J*_SERCA_ is the current density through SERCA (Eq 21), *J*_CaERL_ is the leak current density from ER (Eq 20) and *J*_*D*Cai_ is the intracellular Ca^2+^ diffusion current density between the compartment and its neighboring compartments (Eq 27). The factor *Ar*_ER_ represents the area of ER in the compartment.

The extracellular Ca^2+^ concentration is given by:

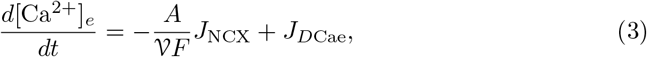

where *J*_*D*Cae_ is the extracellular Ca^2+^ diffusion current density.

The ER Ca^2+^ concentration is governed by the equation [9, 14]:

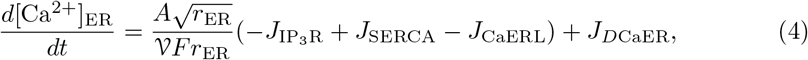

where 𝒱*r*_ER_ represents the volume of ER in the compartment and *J*_*D*CaER_ is the Ca^2+^ diffusion current density between the compartment’s ER and the ERs in neighboring compartments.

The intracellular and extracellular Na^+^ concentrations are modeled as:

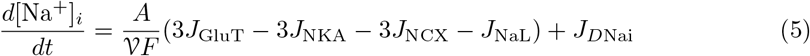

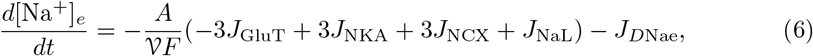

where *J*_GluT_ is the current density through the glutamate transporters (Eq 22), *J*_NKA_ is the current density generate by the transport of Na^+^ through Na^+^/K^+^-ATPase pumps (Eq 23), *J*_NaL_ is the leak current density through the Na^+^ channels (Eq 25), *J*_*D*Nai_ is the intracellular Na^+^ diffusion current density between the compartment and its neighboring compartments and *J*_*D*Nae_ represents the Na^+^ diffusion current density in extracellular space. The factor 3 multiplying the current densities *J*_GluT_, *J*_NKA_ and *J*_NCX_ represents the number of ions carried by these currents [10].

The intracellular and extracellular K^+^ concentrations are modeled according to:

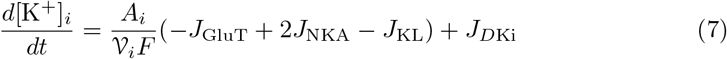

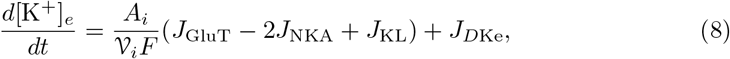

where *J*_KL_ is the leak current density through the K^+^ channels (Eq 26), *J*_*D*Ki_ represents the intracellular K^+^ diffusion current density between the compartment and its neighboring compartments and *J*_*D*Ke_ is the K^+^ diffusion in extracellular space. The factor 2 multiplying the *J*_NKA_ represents the number of K^+^ ions pumped by NKA per cycle [10].

The dependence of the compartment potential *V* on the ionic current densities is described by the following equation:

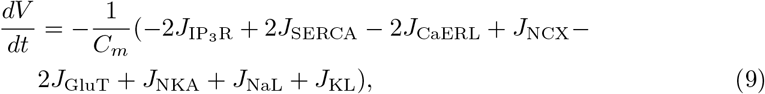

where the integer factors multiplying each current density represent the net current or the number of ions carried per cycle [10].

The intracellular IP_3_ concentration is governed by the equation [9, 14]:

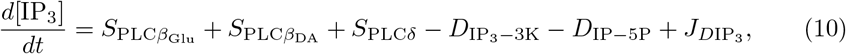

where 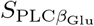 represents the synthesis of IP_3_ by the activation of mGluR (Eq 11), 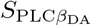 is the synthesis of IP_3_ by the activation of D_1_ receptors (Eq 12), *S*_PLC*δ*_ is the synthesis of IP_3_ by PLC*δ* (Eq 13), 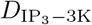 is the degradation of IP_3_ by the IP_3_-3K (Eq 14), *D*_IP−5P_ is the degradation of IP_3_ by IP-5P (equation 15), and 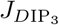 is the intracellular IP_3_ diffusion current density between the compartment and its neighboring compartments.

The production of IP_3_ by the activation of mGluR is related to the extracellular glutamate concentration by the equation [9, 14]:

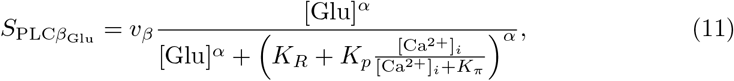

where *v*_*β*_ = 0.674 μM/s is the maximum rate of IP_3_ synthesis by PLCβ, [Glu] is the extracellular glutamate concentration (Eq 16), *α* = 0.7 is the Hill coefficient, *K*_*R*_ = 1.3 μM is the affinity of glutamate to mGluR, *K*_*p*_ = 10 μM is the Ca^2+^-dependent inhibition of PLC and *K*_*π*_ = 0.6 μM is the affinity constant of Ca^2+^ to PLC.

The synthesis of IP_3_ by D_1_ receptors and its dependence on dopamine concentration is modeled as:

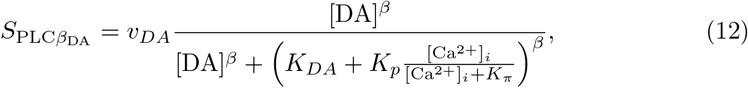

where *v*_*DA*_ = 0.025 μM/s is the maximum production rate of IP_3_ by PLC due to activation of D_1_ receptors, [DA] is the extracellular dopamine concentration (Eq 17), *β* = 0.5 is the Hill coefficient, and *K*_*DA*_ = 5 μM is the affinity of dopamine to the D_1_ receptors. Since the D_1_ receptors also influence the intracellular Ca^2+^ concentration by the activation of the IP_3_ pathway, the dynamics of IP_3_ production by dopamine was modeled similarly to the glutamate-dependent IP_3_ synthesis [9, 11]. The values of the parameters in equation (12) were adjusted in order to reproduce the findings reported in Liu et al [11].

The production of IP_3_ by PLCδ is given by [9, 14]:

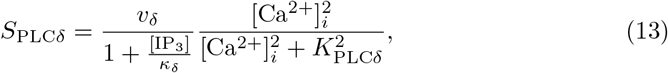

where *v*_*δ*_ = 0.025 μM is the maximum rate of IP_3_ production by PLCδ, *κ*_*δ*_ = 1.5 μM is the inhibition constant of PLCδ and *K*_*P* *LCδ*_ = 0.1 μM is the Ca^2+^-dependent inhibition of PLCδ.

The degradation rates of IP_3_ by IP_3_-3K and IP-5P are modeled according to the equations [9, 14]:

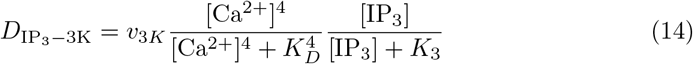

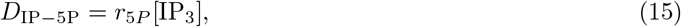

where *v*_3*K*_ = 2 μM is the maximum degradation rate of IP_3_ by IP_3_-3K, *K*_*D*_ = 0.7 μM is the affinity of Ca^2+^ to IP_3_-3K, *K*_3_ = 1 μM is the affinity of IP_3_ to IP_3_-3K and *r*_5*P*_ = 0.04 s^-^1 is the maximum degradation rate of IP_3_ by IP-5P.

The extracellular concentrations of glutamate and dopamine are modeled as:

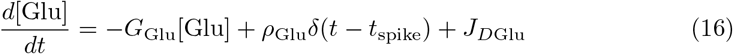

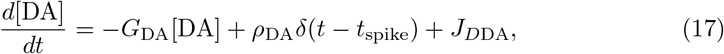

where *G*_Glu_ = 100 s^-1^ and *G*_DA_ = 4.201 s^-1^ are, respectively, the inverses of the time constants of glutamate and dopamine decay, *ρ*_Glu_ = 0.5 μM and *ρ*_DA_ = 3 μM are the amounts of glutamate and dopamine released in each neuron spike, t_spike_ is the presynaptic spike time, and *J*_*D*Glu_ and *J*_*D*DA_ are the glutamate and dopamine diffusion currents in the extracellular space, respectively.

The presynaptic spike times *t*_*spike*_ were drawn from a Poisson distribution with the frequency parameter *f* (in Hz). Since the glutamate that affects the astrocyte activity is released by synaptic terminals ensheathed by astrocytic processes, the glutamatergic stimuli were applied in most of the tests to the distal compartments. This was done to model a tripartite synapse composed by pre- and postsynaptic neurons and the astrocytic process. In some tests, we also applied the glutamatergic stimulation to the somatic compartment. In contrast, since dopamine acts in a diffuse way (volume-transmission), the dopaminergic stimulation was applied to all compartments. More details about the stimulation protocols can be found in the Section “Stimulation Protocols”.

The Ca^2+^ current density through ER by the activation of IP_3_ receptors is given by the following equations:

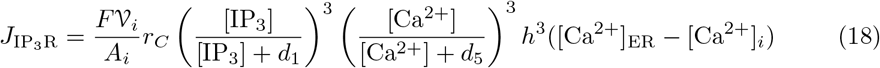

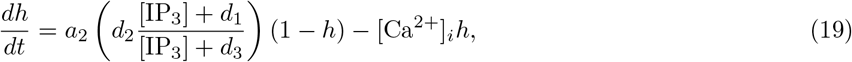

where *r*_*C*_ = 6 s^-1^ is the maximum rate of the Ca^2+^ current through the IP_3_R channel,

*d*_1_ = 0.13 μM and *d*_5_ = 0.08234 μM are, respectively, the dissociation constants of IP_3_ and Ca^2+^, *h* represents the proportion of open IP_3_R channels, *a*_2_ = 0.2 μM^-1^ s^-1^ is the Ca^2+^ inhibition constant, *d*_2_ = 1.049 μM is the Ca^2+^ dissociation constant, *d*_3_ = 0.9434 μM is the receptor dissociation constant of IP_3_ and *d*_1_ = 0.13 μM is the IP_3_ dissociation constant. The biophysical interpretation of these parameters can be found in Ullah *et al*. [12] and Li and Rinzel [13].

The Ca^2+^ leak current density from ER and the SERCA pump of Ca^2+^ into ER are described by the equations:

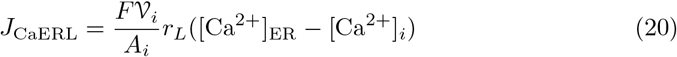

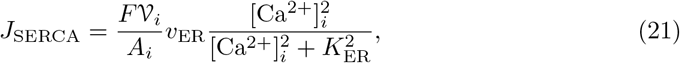

where *r*_*L*_ = 0.11 s^-1^ is the Ca^2+^ leak rate constant, *v*_*ER*_ = 11.93 μM/s is the maximum Ca^2+^ uptake rate by SERCA and *K*_*ER*_ = 0.1 μM is the Ca^2+^ affinity for the SERCA pump.

The ionic current density through GluT is governed by the equation:

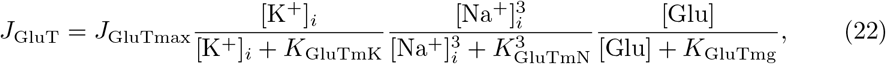

where *J*_GluTmax_ = 0.68 pA/μm^2^ is the maximum transport rate and *K*_GluTmK_ = 5,000 μM/m^2^, *K*_GluTmN_ = 15,000 μM/m^2^ and *K*_GluTmg_ = 34 μM/m^2^ are the half-saturation constants of K^+^, Na^+^ and glutamate, respectively.

The transport of Na^+^ and K^+^ by the Na^+^/K^+^-ATPase pump is modeled as:

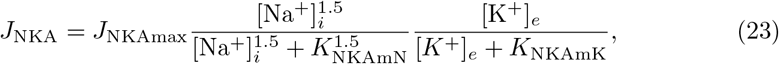

where the maximum pump activity is *J*_NKAmax_ = 1.52 pA/μm^2^, *K*_NKAmN_ = 10,000 μM is the half-saturation constant of Na^+^ and *K*_NKAmK_ = 1,500 μM is the half-saturation constant of K^+^.

The activity of the Na^+^/Ca^2+^-exchanger is given by the equation:

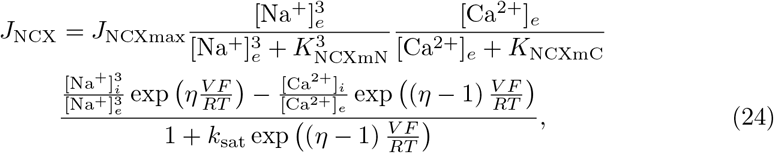

where *J*_NCXmax_ = 0.0001 pA/μm^2^ is the maximum NCX activity, *K*_NCXmN_ = 87,500 μM and *K*_NCXmC_ = 1,380 μM are, respectively, the half-saturation constants of Na^+^ and Ca^2+^, *η* = 0.35 is the energy barrier that controls the voltage dependence of NCX, *R* = 8.314 J /molK is the gas constant, *T* = 303.16 K is the temperature and *k*_sat_ = 0.1 is the saturation constant. In order to ensure that the glutamatergic stimulation of the distal compartments triggers Ca^2+^ signals in the distal, intermediate and proximal compartments, we fixed 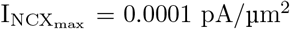. The test carried to determine this value is presented in the Supplementary Materials (Fig. S2)

The Na^+^ and K^+^ leak current densities are governed by the following equations:

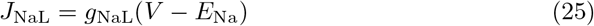

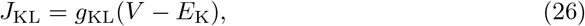

where *g*_NaL_ = 13.482808 S/m^2^ and *g*_KL_ = 145.814171 S/m^2^ are the Na^+^ and K^+^ conductances, respectively, and *E*_Na_ = 61 mV and *E*_K_ = − 94 mV are their corresponding reverse potentials.

All diffusion current densities (linking neighboring intra-, extracellular and ER compartments) are calculated with the following equation [14]:

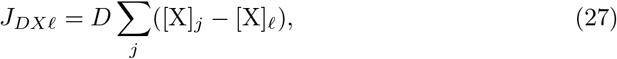

where ℓ is the index of the compartment from/to which the ion/molecule X (in moles) is diffusing, *j* is the index of the compartments connected to the compartment ℓ (for example, ℓ − 1 and ℓ + 1), and *D* is the diffusion coefficient (in s^-^1) of the ion/molecule X. The diffusion current densities *J*_*D*Cai_ (Eq 2), *J*_*D*Cae_ (Eq 3), *J*_*D*CaER_ (Eq 4), *J*_*D*Nai_ (Eq 5), *J*_*D*Nae_ (Eq 6), *J*_*D*Ki_ (Eq 7), *J*_*D*Ke_ (Eq 8), 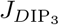 (Eq 10), *J*_*D*Glu_ (Eq 16) and *J*_*D*DA_ (Eq 17) are modeled according to equation (27). The diffusion coefficients of all ions and molecules are given in Table 1 of the Supplementary Materials.

For convenience, all model parameters are also described in Table 1 of the Supplementary Materials. The initial values of [IP_3_], *h* and [Ca^2+^]_ER_ were calculated in order to ensure the equilibrium at the beginning of the simulation and without glutamatergic and dopaminergic stimulation. Similarly, the Na^+^ and K^+^ conductances (*g*_NaL_ and *g*_KL_) were calculated in order to impose equilibrium at the beginning.

When the [Ca^2+^]_*i*_ exceeds the threshold [Ca^2+^]_thresh_ = 0.15 μM [14] we assume there is a Ca^2+^ signal.

### Astrocyte Morphology

We used three different astrocyte morphologies to compare their response and integrative properties. The first morphology consists of a soma with a single branch emanating from it; the second consists of two branches departing directly from the soma; and the third consists of a branch that emanates from the soma and bifurcates farther down into two branchlets. In all three cases, the somatic compartment is a sphere and the remaining compartments are cylinders. The sizes are the same for the three morphologies: the soma radius is 20 μm, the length of the cylindrical compartments is 1 μm and their radii vary from 2 μm to 0.0625 μm (see Fig. 2).

**Fig 2.**
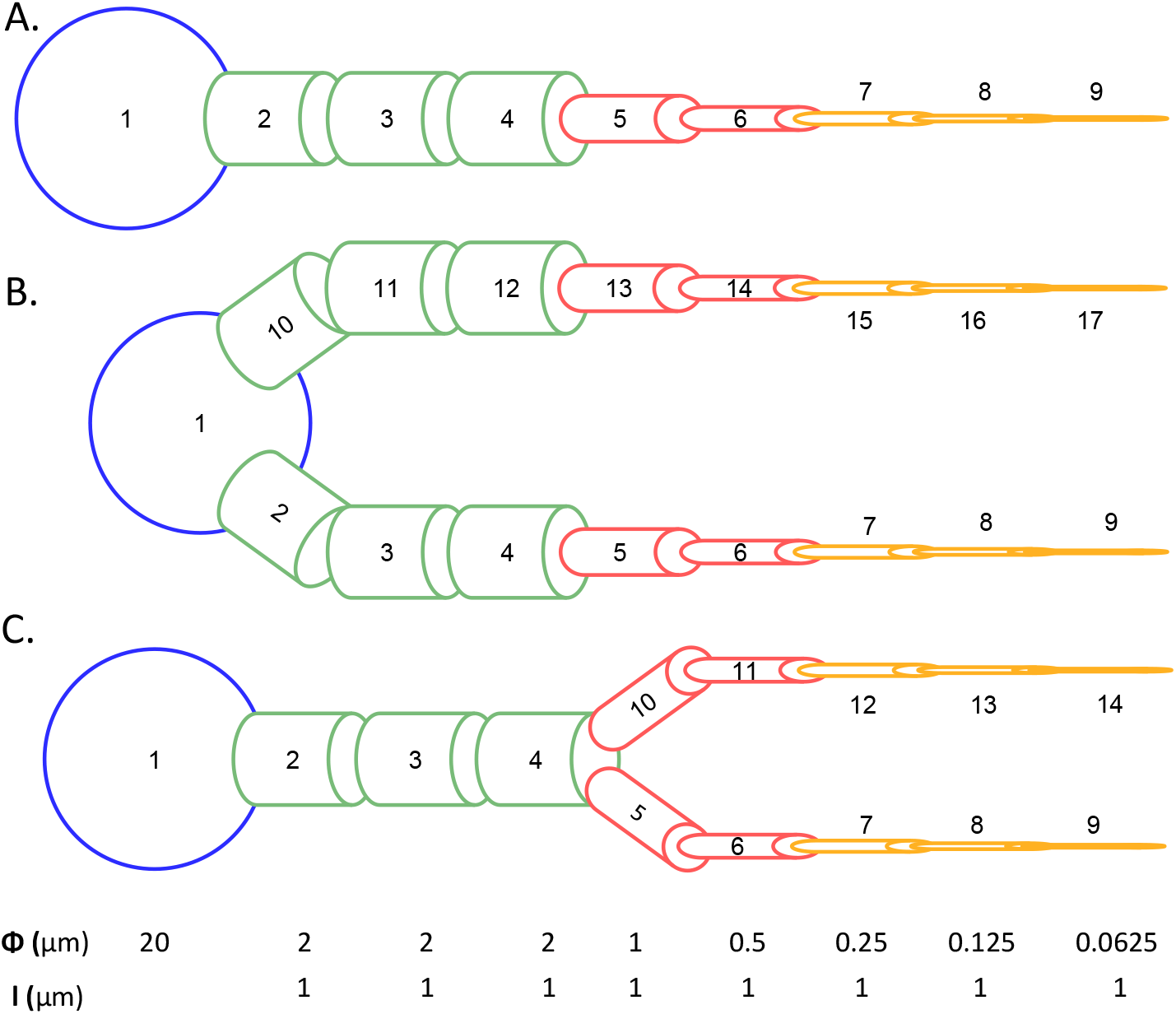
Morphologies of the astrocyte compartmental models. **A**. Linear morphology. **B**. Branched morphology. **C**. Bifurcated morphology. The compartments are numbered in crescent order from soma to distal compartment. The sizes of the soma (blue), proximal (green), intermediate (red) and distal (yellow) compartments are the same for all three morphologies, and are indicated at the bottom as Φ (radius) and I (length). For the branched and bifurcated morphologies, we refer to the branch at the bottom of each cell in the figure as the first branch, and to the branch at the top as the second branch.

### Stimulation Protocols

In all tests, compartments under glutamatergic or dopaminergic stimulation received independent presynaptic inputs modeled as Poisson spike trains with frequency *f* (in Hz) for 100 s. We used two measures to characterize the astrocyte response: (1) the amplitude of the Ca^2+^ signal, which we define as the value of the [Ca^2+^]_*i*_ peak minus the baseline Ca^2+^ concentration, assumed as 0.073 μM; and (2) the number of Ca^2+^ signals triggered by a stimulation.

We organized the tests in order to: a) evaluate and confirm the model response to glutamate and dopamine; b) investigate whether the stimulation of the soma with glutamate or the entire astrocyte with dopamine would enhance the astrocytic response to glutamatergic synaptic inputs arriving at the distal compartments; and c) study whether a glutamatergic synaptic input arriving at one astrocyte branch would modulate the response to glutamatergic synaptic inputs arriving at the other branch. The tests are described below.

#### Glu or DA stimuli: linear morphology

First, we evaluated the response of the model with linear morphology to either glutamatergic or dopaminergic stimulation. In the glutamatergic tests, we considered five different spatial patterns of stimulation: i) glutamatergic input arriving only at the soma (compartment 1); ii) glutamatergic input arriving only at the proximal compartment 3; iii) glutamatergic input arriving only at the intermediate compartment 6; iv) glutamatergic input arriving only at the distal compartment 9; and v) glutamatergic inputs arriving at distal compartments 7,8 and 9. The stimulation frequencies applied were 1 Hz, 5 Hz and 10 Hz. Hence, in total we simulated 15 trials, one for each combination of spatial pattern and frequency. In the dopaminergic tests, all compartments received dopaminergic input. We simulated four trials with dopaminergic stimulation frequencies of 0.1 Hz, 0.5 Hz, 1 Hz and 5 Hz.

#### Integration of Glu stimuli: linear morphology

To test the integrative properties of the linear morphology model to glutamatergic stimulation at different cell positions, we stimulated the soma with glutamate with a frequency *f*_*s*_ and the distal compartments 7, 8 and 9 with a frequency *f*_*d*_. Thirteen frequency values were used for *f*_*s*_ and *f*_*d*_: 0, 0.005, 0.01, 0.05, 0.1, 0.5, 0.75, 1, 2, 3, 4, 5 and 10 Hz. So, in total there were 169 independent trials, one for each possible frequency combination.

#### Integration of Glu and DA stimuli: linear morphology

In these tests, the entire astrocyte was stimulated with dopamine and the distal compartments (7, 8 and 9) were stimulated with glutamate. We used the same 13 frequency values of the previous protocol. Denoting the frequency combination by (*f*_DA_, *f*_*d*_), where *f*_DA_ is the frequency of dopaminergic stimulation and *f*_*d*_ is the frequency of glutamatergic stimulation, there were 169 independent trials, one for each combination.

#### Integration of Glu stimuli: branched/bifurcated morphology

We tested the integrative properties of the branched and bifurcated morphologies to glutamatergic stimulation at different places by co-stimulating the soma with frequency *f*_*s*_, the distal compartments (7, 8 and 9) of the first branch with frequency *f*_*d*_, and the distal compartments (15, 16 and 17 in the branched morphology; 12, 13 and 14 in bifurcated morphology) of the second branch with a fixed frequency of 1 Hz. Thirteen values were used for *f*_*s*_, namely 0, 0.005, 0.01, 0.05, 0.1, 0.5, 0.75, 1, 2, 3, 4, 5 and 10 Hz, and 17 values were used for *f*_*d*_, namely 0, 0.005, 0.01, 0.05, 0.1, 0.5, 0.75, 1, 2, 3, 4, 5, 6, 7, 8, 9 and 10 Hz. Hence, in this protocol we simulated 221 independent trials, one for each frequency combination.

#### Integration of Glu and DA stimuli: branched/bifurcated morphology

To study the effect of simultaneous dopaminergic and glutamatergic stimulation of the models with branched and bifurcated morphologies, we used a protocol that is based on the previous one. In this protocol, all compartments received dopaminergic stimulation with frequency *f*_DA_. We used 13 values for *f*_DA_, namely 0, 0.005, 0.01, 0.05, 0.1, 0.5, 0.75, 1, 2, 3, 4, 5 and 10 Hz. The distal compartments (7, 8 and 9) of the first branch were stimulated with glutamate with frequency *f*_*d*_ and the distal compartments (15, 16 and 17 in the branched morphology; 12, 13 and 14 in the bifurcated morphology) of the second branch were stimulated with glutamate with a fixed frequency of 1 Hz. We used 17 values for *f*_*d*_, namely 0, 0.005, 0.01, 0.05, 0.1, 0.5, 0.75, 1, 2, 3, 4, 5, 6, 7, 8, 9 and 10 Hz. Thus, this protocol comprises 221 independent trials.

### Numerical Methods

The system of equations was numerically integrated using the 4th order Runge-Kutta method with *dt* = 0.01 ms. All numerical routines were implemented in Python 3.7 with the numpy, scipy and numba packages. Since the simulations are time consuming, we used the numba package with JIT compilation and GPU to speed up the simulations. In addition, to save computer memory, we undersampled the time points during the simulation. All codes are available at GitHub (por o endereço).

## Results

We first tested the response of the astrocyte model to glutamatergic or dopaminergic stimulation at different locations and frequencies. In these tests we used only the linear morphology. We will refer to these tests as ‘Glu or DA stimuli: linear morphology’ (see Stimulation Protocols in Methods). Next, we evaluated if somatic glutamatergic stimulation would enhance the response to glutamatergic synaptic inputs arriving at the distal portion of the model with linear morphology. These experiments will be called ‘Integration of Glu stimuli: linear morphology’ (see Stimulation Protocols in Methods). We then evaluated whether the dopaminergic stimulation of the entire astrocyte would have some effect on the behavior observed in the previous experiments. Such experiments will be referred to as ‘Integration of Glu and DA stimuli: linear morphology’ (see Stimulation Protocols in Methods). Subsequently, we evaluated whether the glutamatergic stimulation of one astrocytic branch in the branched or bifurcated morphologies would modulate the response to glutamatergic synaptic inputs arriving at the other branch. These experiments will be called ‘Integration of Glu stimuli: branched/bifurcated morphology’ (see Stimulation Protocols in Methods).

Finally, to study the combined effect of dopamine and glutamate on the behavior of the cell with branched or bifurcated morphology, we repeated experiments of the previous protocol with the addition of dopaminergic stimuli to the whole cell. These experiments will be called ‘Integration of Glu and DA stimuli: branched/bifurcated morphology’ (see Stimulation Protocols in Methods).

In the following subsections, we present the results of our tests organized by the type of experimental protocol as described above.

### Glu stimuli: linear morphology

To study the response of the model with linear morphology to glutamatergic inputs we simulated 15 trials, one for each combination of astrocyte part (soma, compartment 3, compartment 6, compartment 9 or compartments 7,8 and 9) and frequency (1, 5 or 10 Hz).

For glutamatergic input with frequency of 1 Hz, there were only small increases in the [Ca^2+^]_*i*_, below threshold for calcium signal generation, for all stimulation locations (Fig 3, A; first row.). Due to diffusion, stimulation of a given compartment spread passively to all compartments, and the strongest responses occurred for somatic stimulation. Simultaneous stimulation of more compartments produced even stronger responses, as shown for stimulation of the distal compartments.

**Fig 3.**
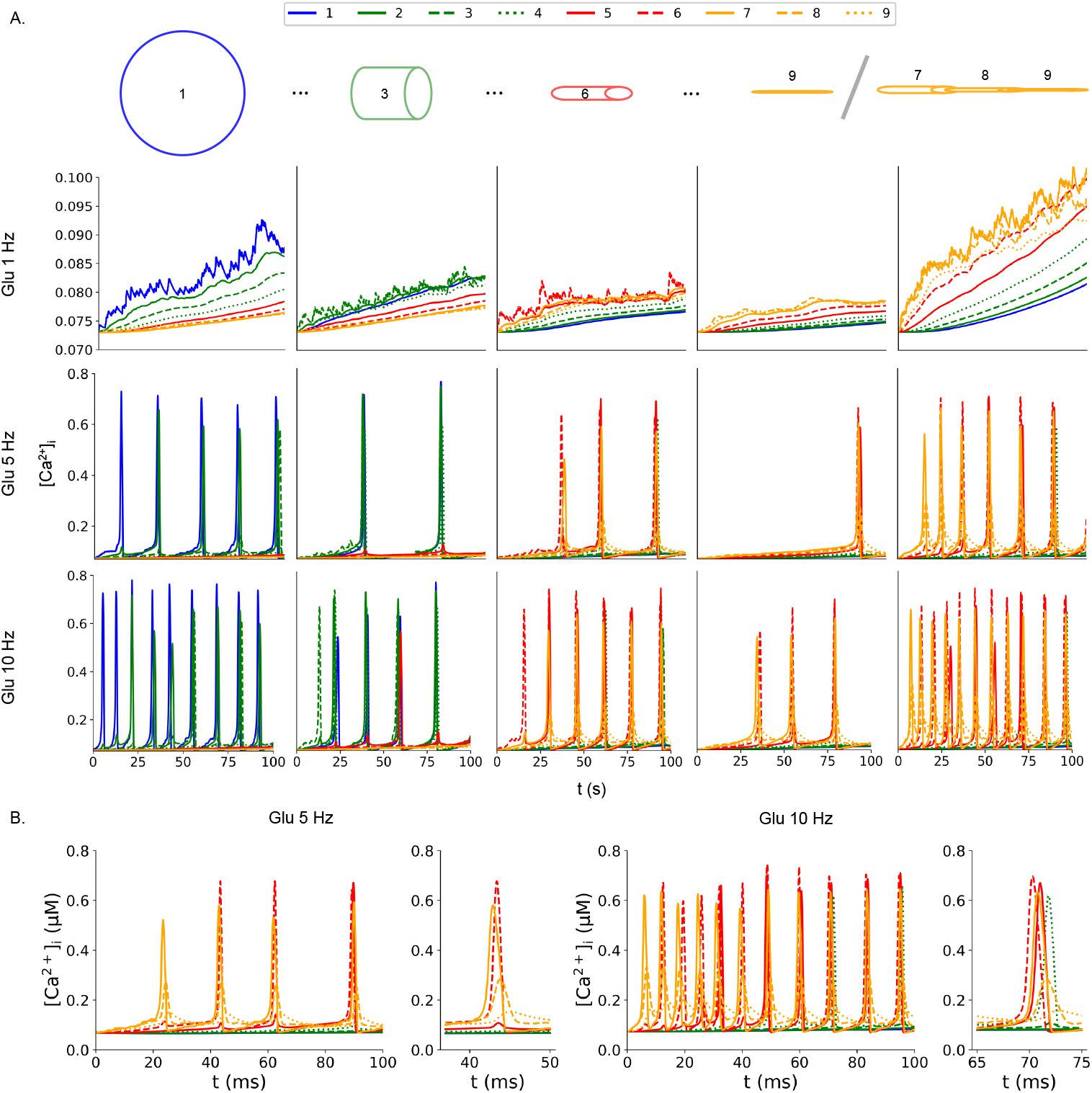
[Ca^2+^]_*i*_ dynamics in the linear morphology with glutamatergic stimulation. Legend indicates the compartment number. **A**. Each column represent a location of glutamatergic stimulation: a) soma in the first column; b) compartment 3 (proximal) in the second one; c) compartment 6 (intermediate) in the third one; d) compartment 9 (distal) in the fourth one; and e) compartments 7, 8 and 9 in the fifth one. Each row indicates the frequency of the glutamatergic stimulation: a) 1 Hz in the first row; b) 5 Hz in the second one; and c) 10 Hz in the third row. **B**. Time course of [Ca^2+^]_*i*_ for trials with glutamatergic input with frequencies of 5 Hz or 10 Hz. For each input frequency, the entire time series is represented in the graphs at the left, amplification of the evoked Ca_2+_ signals are represented in the graphs at the right.

The glutamate stimulation at frequencies of 5 Hz triggered Ca^2+^ signals for any compartment under stimulation (Fig 3, A.). Stimulating only the somatic compartments evoked calcium signals in compartments 1 (soma), 2 and 3 (both proximal). There was a detectable increase in Ca_2+_ concentration of compartment 4. In contrast, stimulation of compartment 3 evoked Ca^2+^ events in compartments 1, 2, 3 and 4, and led to a detectable increase of Ca^2+^ in compartment 5. Stimulation of compartment 6 led to a similar result, evoking Ca^2+^ signals in compartments 4, 5, 6 and 7. Interestingly, no Ca^2+^ event was detected neither in the remaining proximal compartments (2 and 3) nor in the soma. This stimulation also increased Ca^2+^ concentration in compartments 8 and 9. Stimulation of compartment 9 alone evoked Ca^2+^ signals in compartments 5, 6, 7, 8 and 9. Finally, stimulating the compartments 7, 8 and 9 simultaneously triggered Ca^2+^ events in the compartments 4, 5, 6, 7, 8 and 9. Glutamatergic input with frequency of 10 Hz increased the frequency of Ca^2+^ events detected in all compartments for any stimulation point (Fig 3, A.).

Analyzing the time course of [Ca^2+^]_*i*_ with glutamatergic stimulation of 5 Hz, it is possible to note that the Ca^2+^ signals evoked in compartments 6 and 7 showed a spike-like form, while that 8 did not. Similarly, with glutamatergic input with frequency of 10 Hz, the Ca^2+^ events triggered in compartments 4, 5, 6 and 7 are spike-like, while these in compartments 8 and 9 are not. This suggests that the most distal compartment and compartment 3 are affected by the increased [Ca^2+^]_*i*_ that diffused from the compartments in which the Ca^2+^ signals were triggered.

These results confirms that the astrocyte model developed here responds to the stimulation with glutamate, increasing the [Ca^2+^]_*i*_ and evoking Ca^2+^ events. The effect over the astrocyte activity was dependent on the stimulation frequency and compartment under stimulation. For lower synaptic input frequency, there was an increase in the Ca^2+^ concentration without generation of Ca^2+^ signal. Although this low frequency stimulation did not trigger Ca^2+^ signals, it led to an increase in the [Ca^2+^ that diffused to the non-stimulated compartments. In contrast, for higher frequencies (5 Hz and 10 Hz), we detected Ca^2+^ events in the stimulated compartments that propagated toward the non-stimulated ones. The responses detected in the somatic, proximal and intermediate stimulation were stronger when compared to stimulation of the distal compartment. So, these results confirms the model response to glutamatergic stimulation and that the compartments are interconnected by diffusion.

### DA stimuli: linear morphology

In order to simulate the dopaminergic volume transmission [18], all the compartments were stimulated by dopamine with frequencies of 0.1 Hz, 0.5 Hz, 1 Hz and 5 Hz. Similar to the glutamate, all frequencies of dopaminergic stimulation led to an increase in [Ca^2+^]_*i*_ and evoked Ca^2+^ signals (Fig 4, A.). Interestingly, for the stimulation at 0.1 Hz, the signals were detected in the somatic, proximal and intermediate compartments; in the distal portion only in compartment 7 the stimulation evoked a Ca^2+^ signal. With exception of the Ca^2+^ signal evoked in compartment 7, all of them were spike-like. So, the [Ca^2+^]_*i*_ increase in compartment 7 was probably due to diffusion from the intermediate compartments. Increasing the dopaminergic stimulation frequency causes an increase in the number and frequency of the Ca^2+^ signals (Fig 4, B., C. and D.). Higher dopaminergic frequency stimulation also evoked Ca^2+^ signals in the distal compartments.

**Fig 4.**
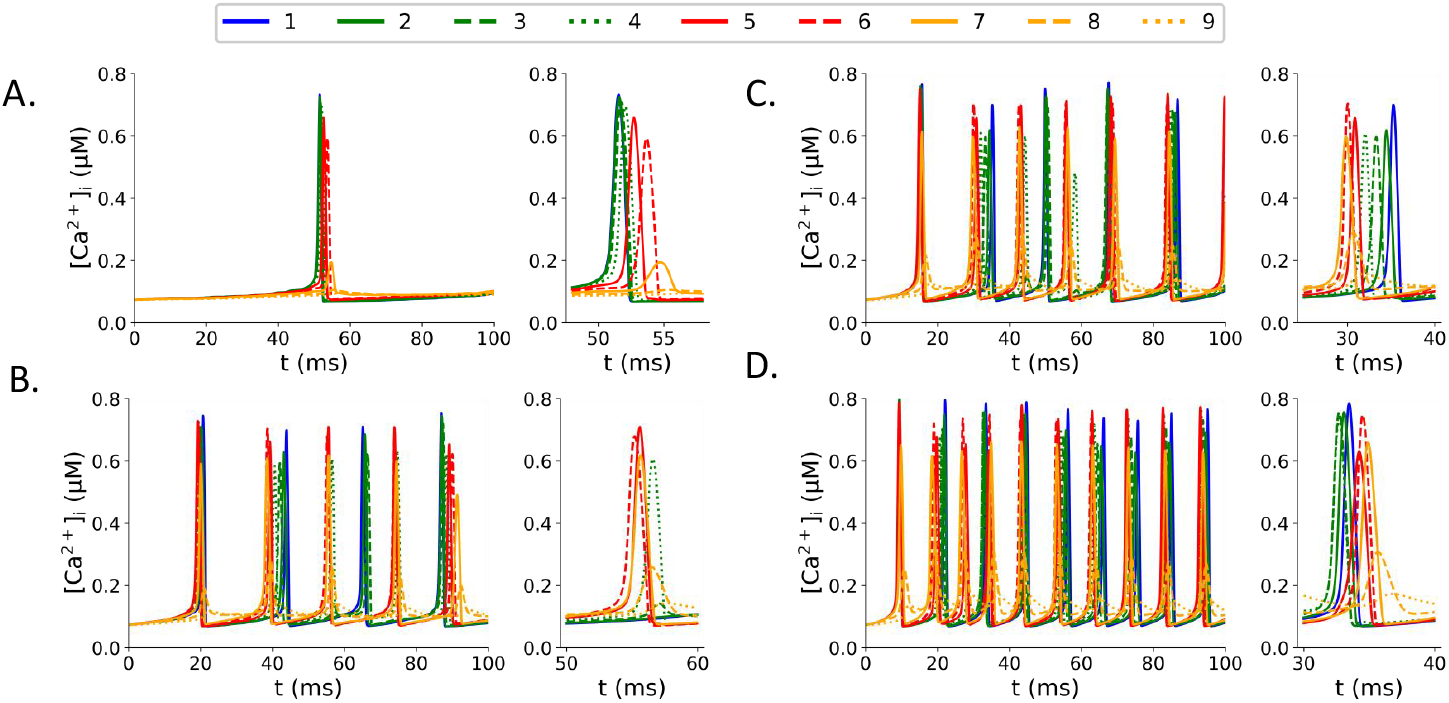
[Ca^2+^]_*i*_ dynamics in the linear morphology with dopaminergic stimulation. The legend indicates the compartment number. A. Dopaminergic stimulation at 0.1 Hz. B. Dopaminergic stimulation at 0.5 Hz. C. Dopaminergic stimulation at 1 Hz. D. Dopaminergic stimulation at 5 Hz.

The response observed confirms the influence of dopamine over the Ca^2+^ dynamics in the present astrocyte model. Since the low frequency stimulation did not trigger Ca^2+^ signals in all distal compartments, the effect observed with higher stimulation frequencies is probably caused by the Ca^2+^ and IP_3_ diffusion from the intermediate and proximal compartments. As the dopamine does not activate the mechanism involving the GluT and the NCX, these results suggest that the glutamate-dependent mechanisms are dominant in the distal portion of the astrocyte process.

### Integration of Glu stimuli: linear morphology

Since the compartments are connected by diffusive currents, there could be an interaction between the stimulation of somatic or proximal compartments with the stimulation of the distal ones by a synaptic input. This stimulation could facilitate and enhance the response to inputs arriving at the distal portion of the astrocyte, increasing the effect of a synaptic terminal over the astrocyte activity. To evaluate this hypothesis, we conducted a trial where both the soma and the distal compartments were stimulated with glutamate. This trial was compared to trials where only the soma or the distal compartments received glutamatergic stimulation.

In the first trial the somatic compartment was stimulated with glutamate with frequency of 5 Hz and the distal compartments 7, 8 and 9 with frequency of 10 Hz. The somatic stimulation led to an increase in the number and frequency of Ca^2+^ signals in all compartments when compared to the stimulation of either the soma alone or the distal compartments alone (Fig 5, C.). Glutamatergic stimulation of the soma alone evoked Ca^2+^ events in the soma and in the proximal compartments 2 and 3, but did not evoke Ca^2+^ signals in the distal part. Stimulation of the compartments 7, 8 and 9 evoked Ca_2+_ signals in all distal and intermediate compartments, but not in somatic and proximal ones (Fig 5, A. and B.)

**Fig 5.**
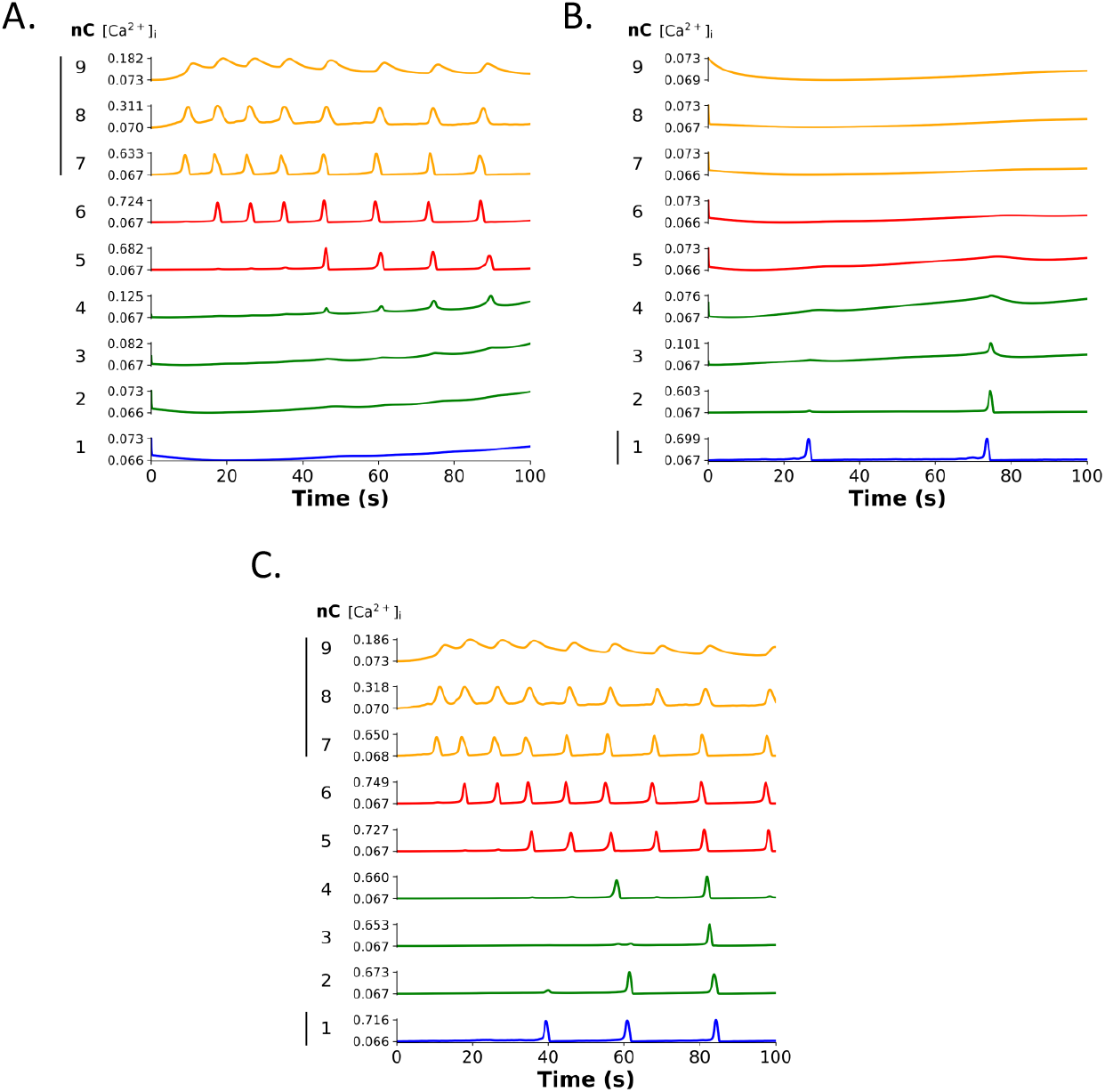
[Ca^2+^]_*i*_ dynamics in the linear morphology with somatic glutamatergic co-stimulation. The black bars indicates the compartments under stimulation. A. Stimulation of the distal compartments (7, 8 and 9) with glutamate with frequency of 10 Hz. B. Stimulation of the soma with glutamate at 5 Hz. C. Stimulation of the soma with glutamate at 5 Hz and distal compartments at 10 Hz.

This result suggests that the somatic stimulation enhanced the effect of the glutamate input to the distal compartments. This facilitation was probably due to the Ca^2+^ and IP_3_ diffusion from the soma to the other compartments. Since neither the stimulation of the soma alone led to the generation of Ca^2+^ signals in the distal compartments nor the stimulation of the distal portion led to Ca^2+^ signals in the soma, the effect observed in the somatic stimulation cannot be attributed to the stimulation of either parts alone. Then, the overall increase in the [Ca^2+^]_*i*_ and the diffusion between compartments facilitated the generation of Ca^2+^ signals in the astrocyte model.

To further explore the interaction between the stimulations of the somatic and distal compartments 7, 8 and 9, we tested different combinations of glutamate stimulation frequencies (*f*_*s*_, *f*_*d*_), where *f*_*s*_ is the somatic frequency and *f*_*d*_ is the distal compartments frequency, and both frequencies in the range from 0 to 10 Hz. The values used for *f*_*s*_ and *f*_*d*_ are given in the Section “Stimulation Protocols”.

Consistent with the previous report, the stimulation of both parts influenced the amplitude and the number of Ca^2+^ signals (Fig 6). In soma, glutamatergic stimulation at the soma was the main factor driving the Ca^2+^ signal generation (Fig 6, A. and D.) However, stimulation of the distal part reduced the necessary frequency at the soma to evoke Ca^2+^ events. Then, for example, stimulating both some and distal compartments with glutamatergic frequency of 0.75 Hz evoked Ca^2+^ signals in the soma. The stimulation of the soma alone or the distal compartments alone with glutamatergic frequency of 0.75 Hz did not trigger Ca^2+^ events in the soma.

**Fig 6.**
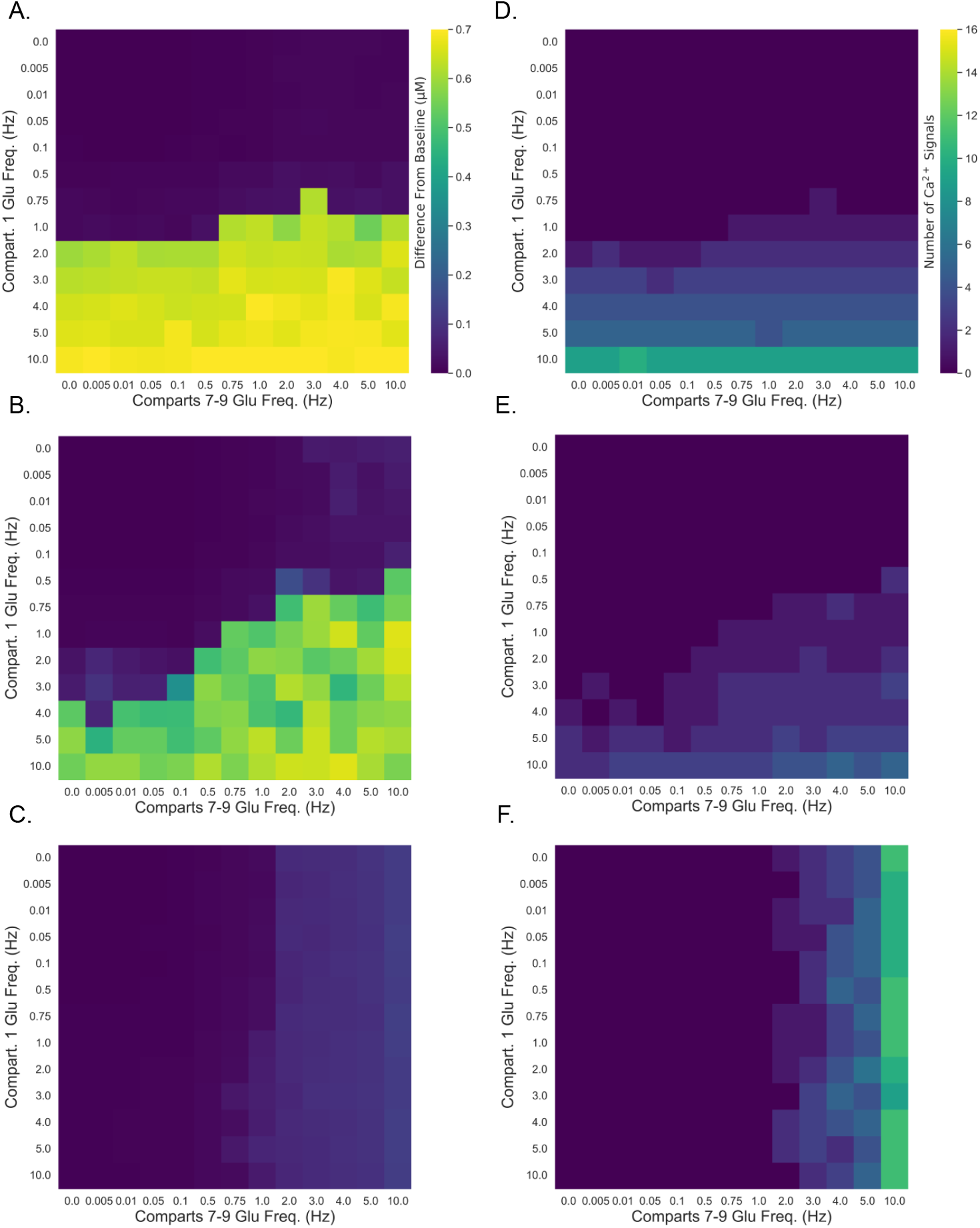
Interaction between the somatic and distal glutamatergic stimulations in the linear morphology. Amplitude of Ca^2+^ signals in A. Soma, B. Compartment 3 (proximal), and C. Compartment 9 (distal). Number of Ca^2+^ signals in the D. Soma, E. Compartment 3 (proximal), and F. Compartment 9 (distal).

A similar pattern was observed in the proximal compartment 3 (Fig 6, B. and E.). Stimulation of the soma evoked Ca^2+^ signals in compartment 3 for frequencies above 4 Hz. In contrast, stimulation of the distal compartments did not evoke Ca^2+^ signals in compartment 3 for any frequency applied. However, combining the stimulation at the soma and in the distal compartments, the frequency needed to evoke these events was reduced. For example, stimulating the soma with glutamatergic frequency of 1 Hz and the distal compartments with frequency of 0.75 Hz evoked Ca^2+^ signals in compartment 3.

Stimulation of the compartments 7, 8 and 9 was the main driving factor in the Ca^2+^ signal generation (Fig 6, C. and F.) Somatic stimulation increase the amplitude of [Ca^2+^]_*i*_ in the compartment 9. However, this stimulation did not increase the number of Ca^2+^ signals in compartment 9, even for the highest frequency applied (10 Hz).

The interaction between the somatic and distal stimulations was more evident in the proximal compartments. Even for lower stimulation frequencies in the distal compartments (that did not elicit Ca^2+^ signals in compartment 3), the co-stimulation with glutamate frequency above 0.1 Hz in the soma facilitated the generation of signals in the proximal compartments. Interestingly, without the stimulation of the distal compartments, the stimulation of the soma alone led to generation of Ca^2+^ signals in compartment 3 for frequencies above 3 Hz. This test showed that combining the stimulation of the soma and the distal compartments facilitated and increased both the amplitude of [Ca^2+^]_*i*_ and the number of Ca^2+^ signals in the somatic and proximal compartments. This effect is probably mediated by diffusion of Ca^2+^ and IP_3_ from the stimulated compartments. Finally, these results suggest that there is a spatial limit for the influence of glutamatergic modulation over the response in the distal compartments. However, this not exclude that a stronger input would modulate the response to glutamatergic synaptic inputs arriving at the astrocyte distal portion.

### Integration of Glu and DA stimuli: linear morphology

To test if the dopamine would enhance the response to glutamatergic synaptic inputs in the distal compartments, we combined the stimulation of the entire astrocyte with dopamine and the astrocyte distal portion with glutamate. Similar to the glutamatergic test, we run simulations for all the combinations of frequencies (*f*_DA_, *f*_*d*_), where *f*_DA_ is the dopaminergic frequency in all compartments and *f*_*d*_ is the frequency in the distal compartments. Both *f*_DA_ and *f*_*d*_ varied from 0 Hz to 10 Hz. More details about the stimulation protocol can be found in the Section “Stimulation Protocols”.

As the glutamatergic test, dopaminergic stimulation also enhanced the response to glutamatergic stimulation of distal compartments (Fig 7). In the soma, the main driving factor evoking Ca^2+^ signals was the dopaminergic stimulation (Fig 7, A. and C.); the stimulation of the distal compartments slightly increased the amplitude of Ca^2+^ signals without changing the number of events detected. Dopaminergic stimulation evoked Ca^2+^ signals in the soma for frequencies above 0.01 Hz, a lower frequency when compared to glutamatergic stimulation. Since the response in proximal and intermediate compartments was similar to the one observed in soma, these data were omitted.

**Fig 7.**
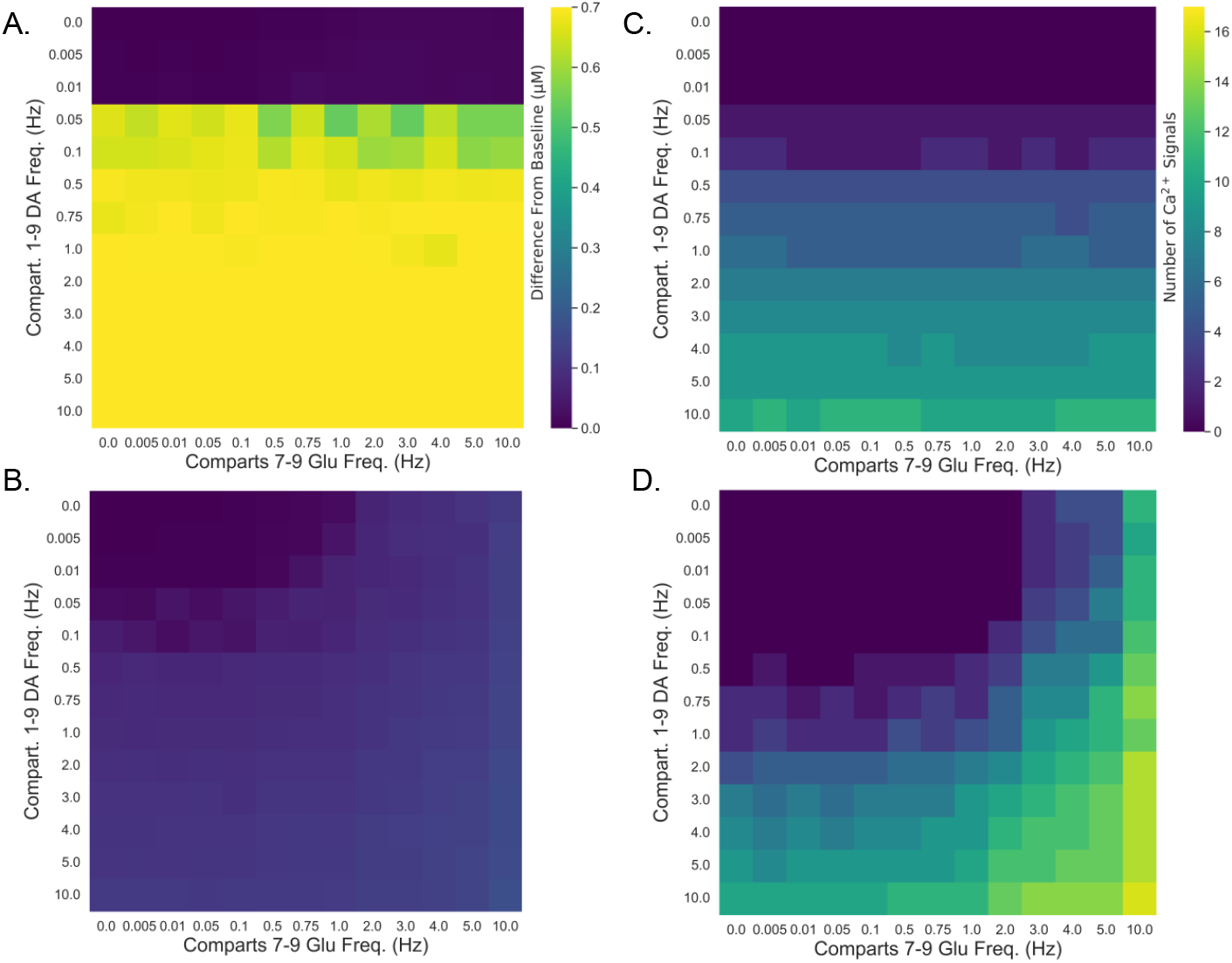
Interaction between the dopaminergic stimulation and glutamatergic inputs at the distal compartments in the linear morphology. Amplitude of Ca^2+^ signals in A. Soma and B. Compartment 9 (distal). Number of Ca^2+^ signals in C. Soma and D. Compartment 9 (distal).

In contrast, in the compartment 9, stimulation with dopamine reduced the frequency necessary to trigger Ca^2+^ events in the distal compartments (Fig 7, B. and D.). Without dopaminergic stimulation, glutamate input generated Ca^2+^ signals for frequencies above 2 Hz. Dopaminergic stimulation with frequency of 0.1 Hz reduced the glutamatergic frequency to 1 Hz; stimulation with 0.5 Hz reduced the frequency to 0.005 Hz (Fig 7, D.) In addition, stimulation of the entire astrocyte with dopamine increased both amplitude and the number of Ca^2+^ events.

These results suggest that dopaminergic stimulation is stronger than the glutamatergic one in the modulation of the response to glutamatergic inputs arriving in the distal portion. Although dopaminergic stimulation with frequencies below 0.5 did not trigger Ca^2+^ signals in distal compartments, even for low frequency dopaminergic stimulation we detected facilitation in the response to glutamatergic inputs in the distal compartment and increase of Ca^2+^ signals amplitude and number. Then, these results showed that dopamine can enhance the response of glutamatergic synaptic inputs in the distal compartments, enabling the glutamate to evoke Ca^2+^ events even for lower synaptic activity.

### Integration of Glu stimuli: branched morphology

In the glutamatergic test, the stimuli applied were combinations (*f*_*s*_, *f*_*d*_) of somatic glutamate stimulation with frequencies *f*_*s*_ varying from 0 Hz to 10 Hz and distal glutamate stimulation (compartments 7, 8 and 9) with frequencies *f*_*d*_ from 0 Hz to 10 Hz (first branch). The compartments 15, 16 and 17 in the second branch received glutamatergic stimulation with frequency of 1 Hz. More details about the stimulation protocol can be found in the Section “Stimulation Protocols”.

Concomitant stimulation of the soma and the distal portion of the first branch facilitated the generation of Ca^2+^ signals in the compartment 15 of the second branch for the low frequency glutamatergic input (Fig 8). Glutamatergic stimulation in the second branch with frequency of 1 Hz did not trigger Ca^2+^ signals by itself.

**Fig 8.**
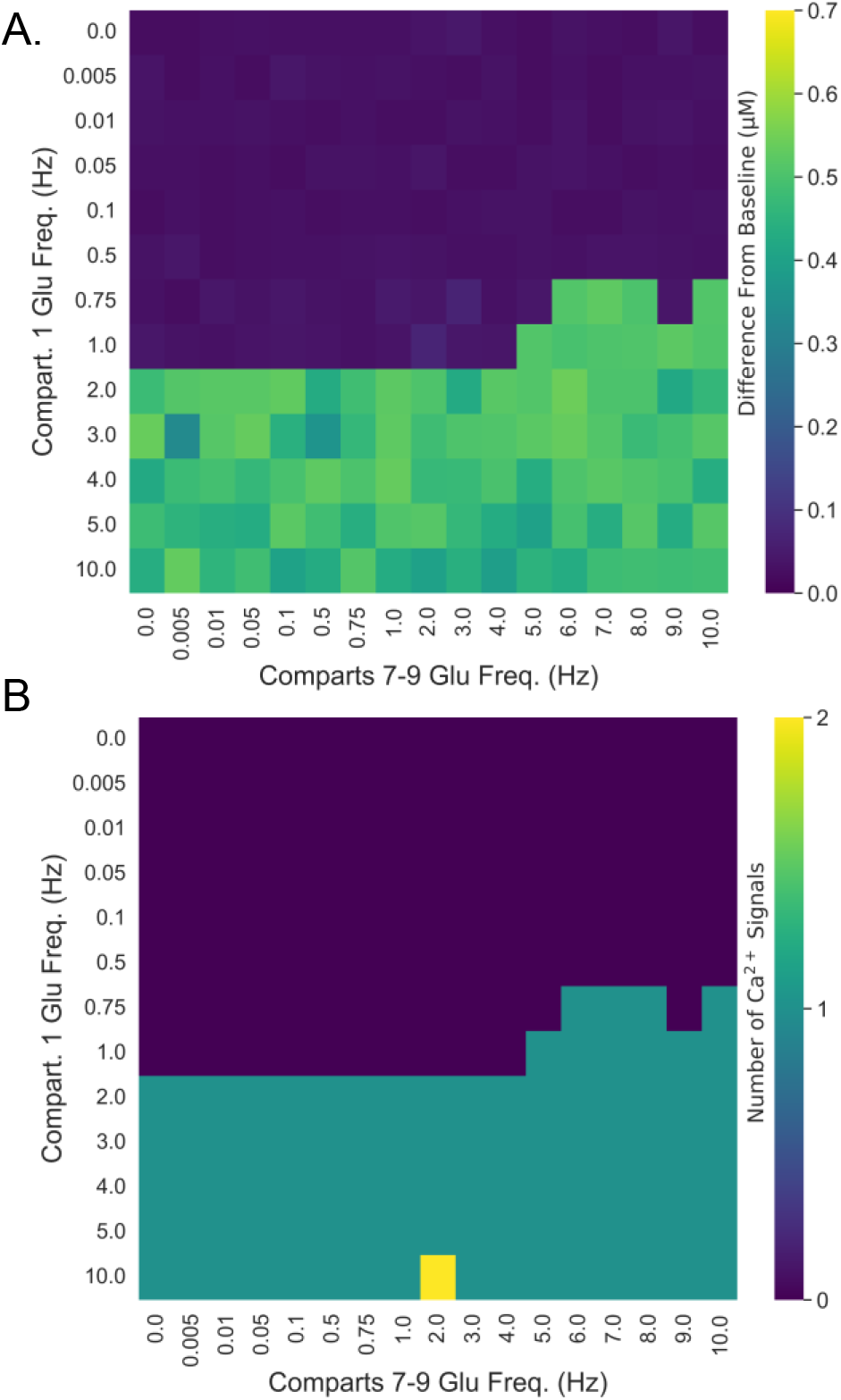
Effect of the co-stimulation with glutamate at the somatic and first branch over the response in the second branch in the branched morphology. Amplitude (A.) and number (B.) of Ca^2+^ signals in compartment 15.

Concomitant stimulation of the first branch did not alter the response in the second branch to the low frequency glutamatergic input. However, stimulation of the somatic compartment with frequencies above 2 Hz facilitated the response in the second branch. This frequency was further reduced by adding the first branch stimulation. Thus, for example, stimulating the soma with glutamatergic input with frequency of 0.75 Hz and the first branch with frequency of 6 Hz evoked Ca^2+^ events in the compartment 15. Nevertheless, there was no Ca^2+^ events in the compartment 17 (most distal part in the second branch) for any combination of stimulations (data not shown).

Then, consistent with the test in the linear morphology, these results show that the somatic activity can enhance the response in the second branch to a low frequency glutamatergic input. This response is further enhanced by stimulating the first astrocyte branch with glutamate. So, the activity in the astrocyte soma can modulate the response in the distal portions and enables the communication between two astrocyte branches that would otherwise function as independent processes. As the tests with the linear morphology suggest, this communication is probably mediated by diffusion of Ca^2+^ and IP_3_ between the compartments. However, since no modulation was observed in the most distal compartment of the second branch, there must be a spatial limit for the communication between two branches of an astrocyte. Consistent with this observation, the stimulation of the first branch without somatic stimulation did not modulate the activity in the second one.

### Integration of Glu and DA stimuli: branched morphology

In the dopaminergic test with the branched morphology, we simulated trials with all frequency combinations (*f*_DA_, *f*_*d*_), where *f*_DA_ is the dopaminergic stimulation frequency varying from 0 to 10 Hz applied in all compartments and *f*_*d*_ is the glutamatergic frequency varying from 0 to 10 Hz applied in the compartments 7, 8 and 9 in the first branch. The compartments 15, 16 and 17 (second branch) received glutamatergic stimulation with frequency of 1 Hz. More details about the stimulation protocol can be found in the Section “Stimulation Protocols”.

Consistent with the tests conducted in the linear morphology, concomitant stimulation with dopamine also facilitated the response in the second branch for the low frequency glutamatergic input (Fig 9). Stimulation of all compartments with dopamine triggered Ca^2+^ signals for frequencies above 0.005 Hz in compartment 15 and above 0.5 Hz in compartment 17 (Fig 9). Since no Ca^2+^ signals were detected by the glutamatergic stimulation of the second branch alone (Fig 8), the addition of dopamine also facilitated the generation of Ca^2+^ signals in the second branch.

**Fig 9.**
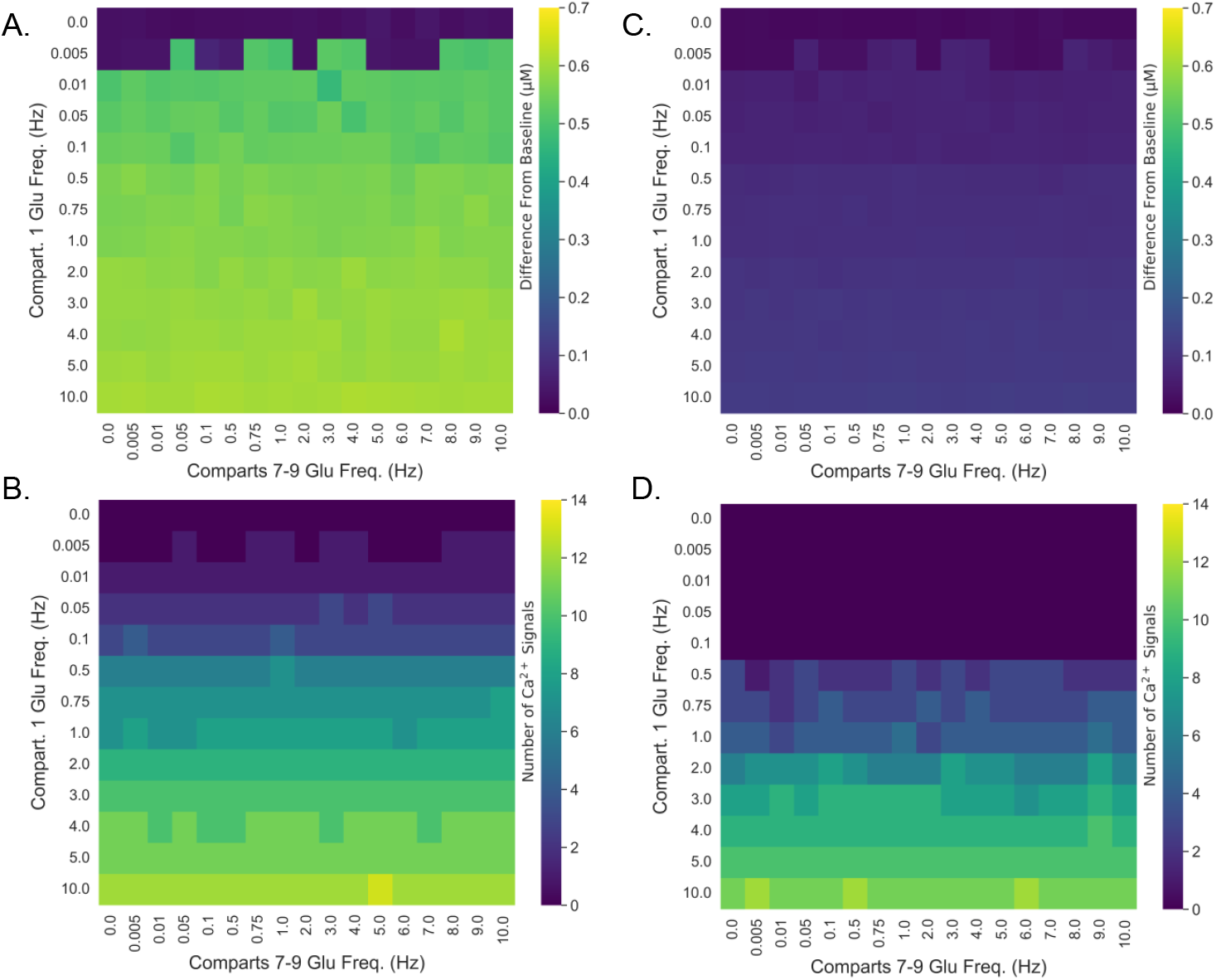
Effect of the co-stimulation with dopamine and first branch with glutamate over the response in the second branch in the branched morphology.. Amplitude of Ca^2+^ signals in the compartments A. 15 and B. 17. Number of Ca^2+^ signals in the compartments C. 15 and D. 17.

However, the glutamatergic input in the first branch did not influence the activity in the second branch. So, the main factors driving the generation of Ca^2+^ signals in the second branch were the dopaminergic stimulation of the entire astrocyte and the low frequency glutamatergic input.

This result suggests that there is a spatial limit for the communication between two astrocytes branches. Since the stimulation of the first branch did not influence the activity of the second one, they acted as independent astrocyte processes. So, inputs arriving in one astrocyte branch would not influence the input processing in other branches. However, with the glutamatergic stimulation of the somatic compartment we detected a facilitation in Ca^2+^ signal generation. So, since dopaminergic input has a stronger effect over the astrocyte activity, we cannot exclude that it could mask the effect of the glutamatergic stimulation in the first branch, which would have a minor effect over the second branch activity.

### Integration of Glu stimuli: bifurcated morphology

In the test with glutamatergic stimulation of the soma, we tested all combinations (*f*_*s*_, *f*_*d*_), where *f*_*s*_ is the stimulation frequency in the soma (from 0 Hz to 10 Hz) and *f*_*d*_ in the frequency in the distal compartments 7, 8 and 9 (first branchlet, from 0 Hz to 10 Hz). The second branchlet received glutamatergic stimulation with frequency of 1 Hz in the distal portion (compartments 12, 13 and 14). More details about the stimulation protocol are described in the Section “Stimulation Protocols”.

The stimulation with glutamate of the first branchlet enhanced the response in the second branchlet to the low frequency glutamatergic input (Fig. 10). There was an increase in both amplitude and number of Ca^2+^ events in compartment 12 by the stimulation of the first branchlet for frequencies above 2 Hz when compared to the stimulation of the second branchlet alone. The somatic stimulation alone facilitated the generation of Ca^2+^ signals in compartment 12 for the frequencies above 4 Hz (Fig. 10). This indicates that somatic activity also modulates the response in the second branchlet.

**Fig 10.**
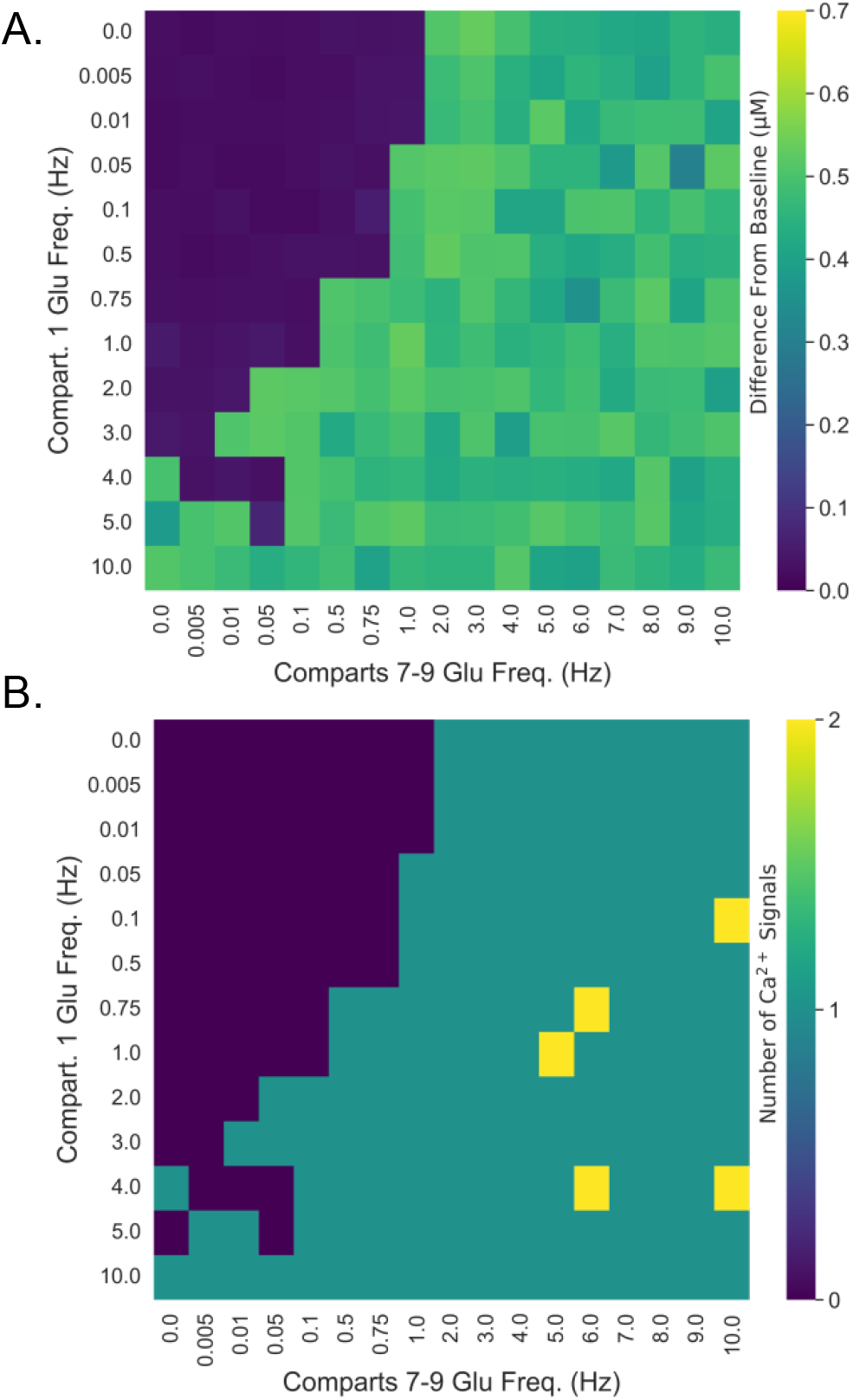
Effects of the somatic glutamatergic co-stimulation in the bifurcated morphology. Amplitude (A.) and number (B.) of Ca^2+^ signal in the compartment 12.

Stimulating the somatic compartment together with the first branchlet also enhanced the response in compartment 12 for the low frequency glutamatergic input (Fig. 10). Addition of somatic stimulation reduced the frequency necessary to detect the influence of the first branchlet stimulation over the second branchlet response. So, for example, applying a stimulation in the soma with frequency of 1 Hz, the stimulation of the first branchlet with frequency of 0.5 Hz evoked Ca^2+^ signals in the second branchlet under the glutamatergic input with frequency of 1 Hz.

However, there was no Ca^2+^ signals in the compartment 14 (most distal one) even for the highest frequency combination tested.

These results indicate that stimulation of either the soma alone or the first branchlet alone enhances the response to a low frequency glutamatergic input in the second branchlet. Interestingly, stimulation of the first branchlet alone with glutamate also modulated the response in the second branchlet, the opposite that was observed in the branched morphology. However, the stimulation of both the soma and the first branchlet interacted, also facilitating the response in the second branchlet. Nonetheless, there is still a spatial limit for this modulation: no interaction was detected in compartment 14 (the most distal one of the second branchlet).

### Integration of Glu and DA stimuli: bifurcated morphology

In the dopaminergic test with the bifurcated morphology, we simulated all frequency combinations (*f*_*DA*_, *f*_*d*_), where *f*_*DA*_ is the dopaminergic input frequency applied to all compartments (from 0 Hz to 10 Hz) and *f*_*d*_ is the frequency of glutamatergic input at the distal portion in the first branchlet (compartments 7, 8 and 9, from 0 Hz to 10 Hz). The second branchlet was stimulated with glutamate at 1 Hz in compartments 12, 13 and 14 in all trials. More details about the stimulation protocol can be found in the Section “Stimulation Protocols”.

Stimulation of the first branchlet with glutamate frequencies above 1 Hz facilitated the generation of Ca_2+_ events in the compartments 12 for the low frequency glutamatergic input (Fig. 11). Similarly, dopaminergic stimulation enhanced the second branchlet response in compartment 12 for frequencies above 0.01 Hz and in compartment 14 above 0.1 Hz. Stimulating all compartments with dopamine reduced the glutamatergic frequency in the first branchlet necessary to enhance the response in the second branchlet. So, for example, dopaminergic stimulation with frequency 0.005 Hz and glutamatergic frequency in the first branchlet to 0.5 Hz together with the glutamatergic stimulation of the second branchlet evoked Ca^2+^ signals in compartment 12.

**Fig 11.**
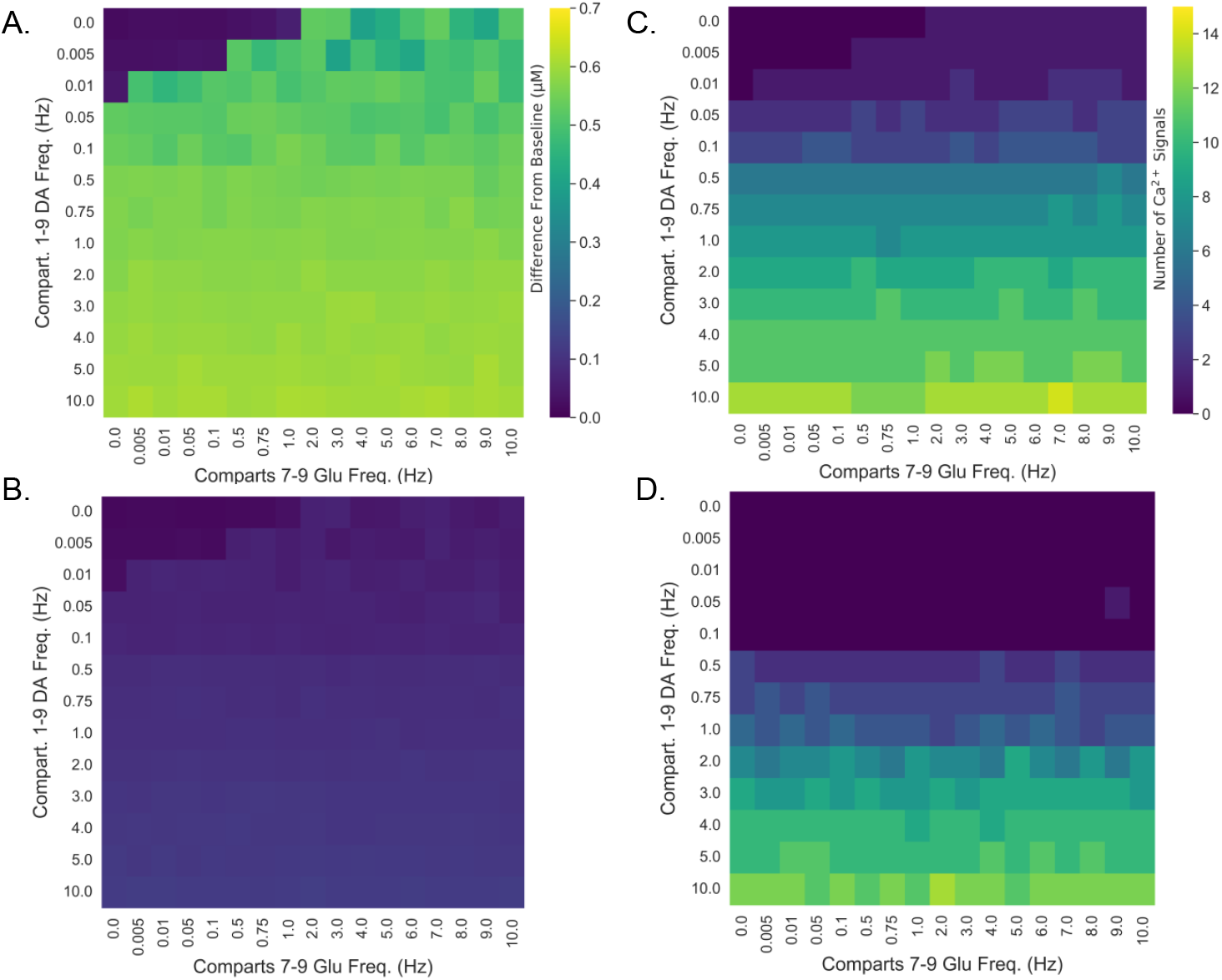
Effects of the dopaminergic co-stimulation in the bifurcated. Amplitude of Ca^2+^ signals in compartment A. 12. and B. 14. Number of Ca^2+^ signals in compartment C. 12 and B. 14.

However, there was no interaction between the dopaminergic input and glutamatergic stimulation in first branchlet with the Ca^2+^ events triggered in compartment 14 (most distal one in the second branchlet). Dopaminergic stimulation was the main driving factor evoking Ca^2+^ signals in this compartment.

This result shows that dopaminergic stimulation and the glutamatergic input arriving at the first branchlet modulate the response in the distal portion of the second branchlet to a low frequency glutamatergic input. In addition, and contrasting with the branched morphology, glutamatergic inputs arriving at the first branchlet enhanced the response in the second branchlet even in the condition without dopaminergic stimulation. However, as seem in the branched morphology, there is a limit for this modulation, since we observed no interaction in compartment 14 response due to dopaminergic stimulation and glutamatergic input in the first branchlet.

## Discussion

In the astrocyte computational model presented here we implemented a mechanism for dopaminergic transmission and its influence over the intracellular Ca^2+^ concentration and the generation of Ca^2+^ signals. We developed an astrocyte compartmental model with three different morphologies, studying the effects of glutamatergic and dopaminergic inputs over the astrocyte activity and its integrative properties. In all morphologies, both glutamatergic and dopaminergic stimulations led to an increase in intracellular Ca^2+^ concentration and triggered Ca^2+^ signals, confirming the model response to these neurotransmitters. Their effects were dependent on the frequency and location of stimulation. Stimulation of the somatic compartment with glutamate or stimulation of the entire astrocyte with dopamine enhanced the response to glutamatergic inputs arriving at the distal portion of the astrocyte in all morphologies. Remarkably, inputs arriving in one process modulated the response to inputs arriving in the second one. This interaction between processes was enabled (branched morphology) or further enhanced (bifurcated morphology) by the glutamatergic stimulation of the soma or dopaminergic stimulation of the entire astrocyte.

The time course of Ca^2+^ dynamics detected in the present work is consistent with other computational models of astrocytes [9, 10] and experimental data [17]. Ca^2+^ signals detected in the soma, proximal and intermediate compartments showed a spike-like form, with a sharp ascending phase and a posterior descending phase. In distal compartments, which had a lower ER-cytosol volume ratio (*r*_ER_), the response had a lower amplitude compared to the other compartments in the model. In addition, the Ca^2+^ signal had a longer duration in distal compartments when compared to the remaining compartments. As shown in the computational model of astrocytes developed by Oschmann *et al*. [10], compartments with smaller radius showed a reduced amplitude and oscillation frequency of glutamate-triggered Ca^2+^ signals. These differences were attributed to the distinct glutamatergic mechanisms regulating Ca^2+^ dynamics in each astrocyte portion. Similarly, we also observed in the present model that varying the parameters of maximum NCX activity and GluT currents affected the Ca^2+^ signal amplitude (S2 Fig.). So, the differences detected between the compartment responses in the present work can also be attributed to variations in the *r*_ER_ and to the different glutamatergic mechanism in each astrocyte region. In smaller compartments, which have lower *r*_ER_ values, the GluT-dependent mechanism has a stronger effect over the generation of Ca^2+^ signals. In contrast, in compartments with greater radius, higher *r*_ER_) values, the dominant mechanism is dependent on the activation of mGluR receptors.

Interestingly, low frequency dopaminergic stimulation did not trigger Ca^2+^ signals in the distal compartments. This corroborates the notion that the GluT-NCX mechanism has a stronger effect over the Ca^2+^ concentration in astrocyte regions with lower *r*_ER_. Since the major glutamatergic input to astrocyte comes from synaptic contact [19], it also suggests that the dopaminergic transmission modulates the astrocyte activity in larger processes and, so, regions that do not receive direct synaptic input. This observation is consistent with the diffuse dopaminergic transmission, affecting larger areas compared to glutamate [18].

Diffusion of Ca^2+^ and IP_3_ between astrocyte regions is a third mechanism that can modulate the intracellular Ca^2+^ concentration and influence the generation of Ca^2+^ signals. In the present work, by stimulating only the distal compartments, we detected an increase in intracellular Ca^2+^ concentration and, sometimes, generation of Ca^2+^ signals in non-stimulated compartments. In some cases, the increased Ca^2+^ concentration caused by diffusion from other compartments reached the threshold for counting a Ca^2+^ signal without the generation of an event with the spike-like form.

This indicates that just by diffusion, it would be possible to detect gliotransmitter release without the detection of a spike-like form Ca^2+^ signal. High frequency dopaminergic stimulation evoked Ca^2+^ events in the astrocyte distal portions in the present work, suggesting that dopamine modulate the response in these compartments by diffusion of Ca^2+^ and IP_3_ from the intermediate, proximal and somatic compartments. Since the gliotransmitter release is dependent upon the increase in intracellular Ca^2+^ concentration, this results suggest that gliotransmitter could be released in astrocyte locations that are not under direct synaptic stimulation, enabling the astrocyte to modulate a broader area and affect other neurons.

Then, diffusion of Ca^2+^ and IP_3_ from other compartments can also facilitate the generation of Ca^2+^ signals by bringing the Ca^2+^ and IP_3_ concentrations closer to the threshold for triggering a Ca^2+^ event. Since during a Ca^2+^ signal the Ca^2+^ and IP_3_ concentrations are increased, it can cause a further diffusion toward other astrocyte regions. This phenomenon of activity spread along astrocytes was named “calcium waves”. In principle, the diffusion could be a mechanism responsible for the generation these Ca^2+^ waves.

However, as showed by Bindocci *et al*. [17], the Ca^2+^ activity in astrocytes is rather a local process, with events triggered in one branch hardly reaching the soma. That is, the astrocyte processes show in general an activity compartmentalization in which the activation of a process alone do not interfere in the activity of another region in the same astrocyte. So, it is unlikely that a signal triggered in the distal portion would travel the astrocytic process and triggers a second Ca^2+^ event in proximal or somatic regions.

Indeed, in the present work, stimulation of compartments 1, 3, 6 and 9 separately did not evoke Ca^2+^ signals in distant compartments. So, for example, stimulating only the compartment 1 (soma) with glutamate, we detected Ca^2+^ events in compartments 2, 3 and 4, even for high frequency stimulation. Similarly, stimulating the compartments 7, 8 and 9 simultaneously evoked Ca^2+^ signals at most in the compartment 4. Thus, the astrocyte model developed here also show that activity compartmentalization.

Nonetheless, Bindocci *et al*. [17] also recorded what they called “global events”. These events were characterized by the activity of several astrocyte regions at the same time. Interestingly, global events were associated with movement in mice. Thus, the astrocytic compartmentalization described above do not exclude the possibility that a stimulation arriving at different points could generate the global events. In addition, since diffusion of Ca^2+^ and IP_3_ can interfere in the activity of astrocytic processes, the release of glutamate in points close to the soma or proximal portions could in principle facilitates the response in other regions and trigger the global events during relevant behaviors. However, as the glutamatergic transmission is more restricted to synaptic terminals [19], the glutamate release is unlikely to induce these global events.

Nonetheless, neuromodulators, such as dopamine, for presenting diffuse transmission that affects large brain areas, are particularly suited to trigger and modulate these global events [18].

Then, a clear characterization of these local and global events is crucial to understand how astrocytes can integrate inputs arriving at different processes or regions and how they can modulate large neural networks [20]. In this sense, to test possible glutamatergic and dopaminergic mechanisms for the generation of global events and their effects over the integrative properties in astrocytes, we evaluated whether the co-stimulation of somatic compartment with glutamate or the stimulation with dopamine of the entire astrocyte would enhance the input integration and facilitated astrocyte response. In the linear morphology, this stimulation protocol together with stimulation at the distal compartments indeed enhanced the response in the proximal and intermediate processes, increasing both amplitude and frequency of Ca^2+^ signals. Hence, a stimulus that would not by itself evoke Ca^2+^ events starts activating other regions in the same astrocyte, characterizing the global events. Therefore, these results suggest that the global events can be triggered by two concomitant types of stimulus: a) a synaptic glutamatergic input arriving in distal portions; b) a modulatory input, affecting somatic or proximal regions. As discussed above, since glutamatergic input are predominantly in synaptic regions [19], it is unlikely that glutamate modulate the somatic and proximal astrocyte activity. So, dopaminergic stimulation is a possible candidate for this modulation.

A possible role for the astrocyte Ca^2+^ global events would be the activity integration in different astrocyte processes. Since this glial cells are involved in modulation of large neuron networks [20], a global event could then be associated with this large modulatory role. In the present work we addressed this question using both branched and bifurcated morphologies. In the branched morphology, stimulating the first branch had no effect over the activity in second branch. However, by stimulating the somatic compartment with glutamate enabled that inputs at the first branch modulated the response of the second branch. Thus, stimulating both the somatic compartment facilitated the integration of inputs arriving at different astrocyte processes and enhanced the response in the branch receiving a weaker input.

A similar result was obtained with the bifurcated morphology. However, in this case, the stimulation of the second branch alone already modulated the response in the second branch. Applying glutamatergic stimulation in the soma or dopaminergic stimulation of the whole cell further enhanced the modulation effect of the first branch stimulation over the second branch activity. So, these results suggest that the communication of different astrocyte branches is limited by spatial constraints: branches that have a closer intersection point have also a stronger influence over each other. In addition, it is necessary that one of the branches receive a high frequency synaptic input in order to effectively modulate the second branch activity. This frequency is lowered by adding the glutamatergic or dopaminergic modulatory input. In this sense, somatic and proximal astrocyte regions can act as an integrator of the processes activity and, when activate by the modulatory inputs, enhance the communication between branches.

The effects observed in these tests can be associated with the diffusion of Ca^2+^ and IP_3_. The stimulation of the somatic compartment and of the first branch in the branched and bifurcated morphologies induce an increase in Ca^2+^ and IP_3_ concentrations. From the points under stimulation, these compounds can diffuse towards the remaining compartments, in particular to another astrocyte branch, also increasing their concentration and then facilitating the response in these compartments. Since diffusion has a spatial limit for its influence, as shown by comparing the branched and bifurcated morphologies, it could explain the spatial constraint detected in the communication between astrocyte branches. Although Bindocci *et al*. [17] did describe a localized astrocyte activity, it is important to note that Ca^2+^ and IP_3_ diffusion could pass undetected in experimental settings, possibly explaining the absence of a diffusion signal and the apparent compartmentalization in astrocytes. Consistent with this finding, mice genetically modified to express a “sponge” that traps IP_3_ show a reduction in the amplitude and frequency of Ca^2+^ signals [21]. So, the integrative enhancement by glutamatergic and dopaminergic modulatory inputs can be attributed to the diffusion of Ca^2+^ and IP_3_ from the astrocyte stimulated regions, induce diffusion to other parts and facilitate the response in several regions, possibly generating a global event.

Release of gliotransmitters from astrocyte is tied to the Ca^2+^ concentration. Since these molecules can modulate synaptic activity, enhancing the communication between two neurons [14], by means of global events, astrocytes could influence several neurons at the same time and modulate their synchronization. Neuromodulators as dopamine, could then be associated with neuron synchronization by activating several astrocyte regions and promoting gliotransmitter release. Indeed, dopaminergic transmission in prefrontal cortex and basal ganglia is associated with modulation of oscillatory activity [25]. Therefore, by triggering global event, modulatory inputs to astrocyte can influence several synapses at the same time, linking relevant behaviors or salient stimuli to wide neuron activity.

However, more studies are needed to elucidate the role of dopamine transmission and its effect over the astrocyte modulation of a neural network. Future works of our group will address this questions: how the dopaminergic transmission can modulate the astrocyte activity and as such interfere in the activity of a neural network.

### Limitations

Some limitations of the present work should be considered in order to correctly interpret the model developed here and the results obtained within. First, we cannot exclude other pathways for dopaminergic modulation over intracellular Ca^2+^ concentration. Indeed, a study found that oxygen reactive species produced by the degradation of dopamine by monoamine oxidase affect the Ca^2+^ dynamics in astrocytes [22]. Dopamine can also modulates astrocyte activity by interacting with the cyclic adenosine pathway, that in turn modulate Na^+^ and K^+^ channels [23]. In addition, dopamine is not the only neuromodulator found to influence astrocyte intracellular Ca^2+^ concentration. Noradrenaline, serotonin and acetylcholine also impact astrocyte activity [20].

The morphologies and their parameters used in the present study is rather a simplification and generalization of common features observed in astrocyte morphology. A typical rodent protoplasmic astrocyte can emit 5 to 10 primary processes from which the peripheral processes emanate [1]. In addition, there are physiological and morphological heterogeneity between astrocytes subtypes and the brain structure in which they are located [1]. However, the computational constrains imposes a limit to the simulation of an astrocyte with detailed morphological architecture and its parameters. Nonetheless, although using an idealized version of astrocytes, the modeling and simulating approaches give several insights about the physiology of astrocytes that can guide future theoretical and experimental studies.

## Conclusion

Here we presented an astrocyte computational model that implements the dopaminergic modulation over the intracellular Ca^2+^ concentration. That modulatory input can enhance the integration between synaptic inputs arriving in different astrocytic processes. Generation of global events by dopamine can be related to these integration effects, enabling the communication between astrocytic branches that would otherwise function independently. This bridge can facilitate the response to low frequency synaptic input and lead to the simultaneous modulation of several synapses. We detected different modulatory effects by comparing three morphologies, showing that these effects are dependent on spatial characteristics. So, the integrative properties in astrocytes are closely linked to the astrocyte morphology. Nonetheless, more studies are needed to test the model predictions presented here and to unveil the entire influence of dopaminergic transmission over the astrocyte activity.

## Supporting information

**S1 Table. Model parameters**.

**S2 Fig. Effect of GluT and NCX currents over the Ca**^**2+**^ **signals in Distal Compartments**. To test the influence of the currents through GluT and NCX over the Ca^2+^ dynamics, we simulated trials with combinations of different values for the parameters *J*_*GluT* *max*_ and *J*_*NCXmax*_. In this test, compartment 9 received glutamatergic synaptic input simulated as a Poisson spike train with frequency of 10 Hz for 100 s. With *J*_*NCXmax*_ = 0 there were an increase in the amplitude of the [Ca^2+^]_*i*_ in distal compartments and an increase in the number of Ca^2+^ signals triggered in the distal compartments when compared to the conditions with higher values of *J*_*NCXmax*_. In compartment 8, NCX current is the a dominant factor regulating [Ca^2+^]_*i*_, as the variation of *J*_GluTmax_ did not change the [Ca^2+^]_*i*_ amplitude. However, in compartment 9, the increase in *J*_GluTmax_ led to a slight increase in the amplitude of [Ca^2+^]_*i*_. Since the number and amplitude of Ca^2+^ are reduced by higher values of NCX current attenuates the effects of glutamatergic stimulation over the [Ca^2+^]_*i*_ and its propagation. A. [Ca^2+^]_*i*_ time series for three values of *J*_*NCXmax*_ (upper: 0; middle 0.0001 pA/μm^2^; bottom: 0.001 pA/μm^2^). B. and C. [Ca^2+^]_*i*_ amplitudes in compartment 8 (B.) and 9 (C.)

## Acknowledgments

T.O.B. is supported by a CAPES PhD scholarship. A.C.R. is supported by a CNPq grant 303235/2018-7. This article was produced as part of the activities of FAPESP Research, Innovation and Dissemination Center for Neuromathematics (Grant No. 2013/07699-0, S. Paulo Research Foundation).

