## Supplementary material for "Dopamine facilitates the response to glutamatergic inputs in a computational model of astrocytes"

### Dopamine Enhances Input Integration in a Compartmental Model of Astrocytes - Supplementary Material

August 2022

Table 1: Model parameters.

| Parameter | Value | Unit |
|---|---|---|
| Resting State | | |
| $[Ca^{2+}]_i$ | 0.073 | $\mu M$ |
| $[Ca^{2+}]_{ER}$ | 21.978 | $\mu M$ |
| $[Ca^{2+}]_e$ | 1800 | $\mu M$ |
| $[Na^+]_i$ | 15000 | $\mu M$ |
| $[Na^+]_e$ | 145000 | $\mu M$ |
| $[K^+]_i$ | 100000 | $\mu M$ |
| $[K^+]_e$ | 3000 | $\mu M$ |
| $[IP_3]$ | 0.1917 | $\mu M$ |
| $h$ | 0.8028 | — |
| $V$ | -85 | mV |
| $[Glu]$ | 0 | $\mu M$ |
| $[DA]$ | 0 | $\mu M$ |
| $IP_3$ Dynamics | | |
| PLC$\beta$ Synthesis | | |
| $K_p$ | 10 | $\mu M$ |
| $K_\pi$ | 0.6 | $\mu M$ |
| PLC$\delta$ Synthesis | | |
| $v_\delta$ | 0.025 | $\mu M/s$ |
| $\kappa_\delta$ | 1.5 | $\mu M$ |
| $K_{PLC\delta}$ | 0.1 | $\mu M$ |
| $IP_3$-3K Degradation | | |
| $v_{3K}$ | 2 | $\mu M/s$ |
| $K_D$ | 0.7 | $\mu M$ |
| $K_3$ | 1 | $\mu M$ |
| IP-5P Degradation | | |
| $r_{5P}$ | 0.04 | $1/s$ |
| Glutamate Transmission | | |

| | | |
|---|---|---|
| $\rho_{glu}$ | 0.5 | $\mu M$ |
| $G_{glu}$ | 100 | 1/s |
| $K_R$ | 1.3 | $\mu M$ |
| $v_\beta$ | 0.647 | $\mu M/s$ |
| $\alpha$ | 0.7 | — |
| Dopamine Transmission | | |
| $\rho_{DA}$ | 3 | $\mu M$ |
| $G_{DA}$ | 4.201 | 1/s |
| $v_{DA}$ | 0.025 | $\mu M/s$ |
| $K_{DA}$ | 5 | $\mu M$ |
| $\beta$ | 0.5 | — |
| ER Leak Current | | |
| $r_L$ | 0.11 | 1/s |
| SERCA Current | | |
| $v_{ER}$ | 11.93 | $\mu M/s$ |
| $K_{ER}$ | 0.1 | $\mu M$ |
| h Dynamics | | |
| $d_1$ | 0.13 | $\mu M$ |
| $d_5$ | 0.08234 | $\mu M$ |
| $d_2$ | 1.049 | $\mu M$ |
| $d_3$ | 0.9434 | $\mu M$ |
| $a_2$ | 0.2 | $1/(\mu M * s)$ |
| $r_C$ | 6 | $1/s$ |
| GluT | | |
| $I_{GluTmax}$ | 0.68 | $pA/\mu m^2$ |
| $K_{GluTmN}$ | 15000 | $\mu M$ |
| $K_{GluTmK}$ | 5000 | $\mu M$ |
| $K_{GluTmg}$ | 34 | $\mu M$ |
| NKA | | |
| $I_{NKAmax}$ | 1.52 | $pA/m^2$ |
| $K_{NKAmN}$ | 10000 | $\mu M$ |
| $K_{NKAmK}$ | 1500 | $\mu M$ |
| NCX | | |
| $I_{NCXmax}$ | 0.001 | $pA/m^2$ |
| $K_{NCXmN}$ | 87.5 | $\mu M$ |
| $K_{NCXmC}$ | 1.380 | $\mu M$ |
| $k_{sat}$ | 0.1 | — |
| $\eta$ | 0.35 | — |
| Voltage Parameter | | |
| $C_m$ | 0.01 | $F/m^2$ |
| Leak Currents | | |
| $g_{Naleak}$ | 13.482808 | $S/m^2$ |

| | | |
|---|---|---|
| $E_{Na}$ | 61 | mV |
| $g_{Kleak}$ | 145.814171 | $S/m^2$ |
| $E_K$ | -94 | mV |
| Diffusion Constants | | |
| $D_{Ca}$ | 0.2 | $1/s$ |
| $D_{CaER}$ | 0.001 | $1/s$ |
| $D_{IP_3}$ | 0.2 | $1/s$ |
| $D_{Na}$ | 0.316 | $1/s$ |
| $D_K$ | 0.938 | $1/s$ |
| $D_{Cao}$ | 4.52 | $1/s$ |
| $D_{Nao}$ | 26.6 | $1/s$ |
| $D_{Ko}$ | 1.732 | $1/s$ |
| $D_{glu}$ | 4e-4 | $1/s$ |
| $D_{DA}$ | 13.8 | $1/s$ |
| Physical Constants | | |
| T | 303.16 | K |
| F | 96500 | $C/mol$ |
| R | 8.314 | $J/mol * K$ |

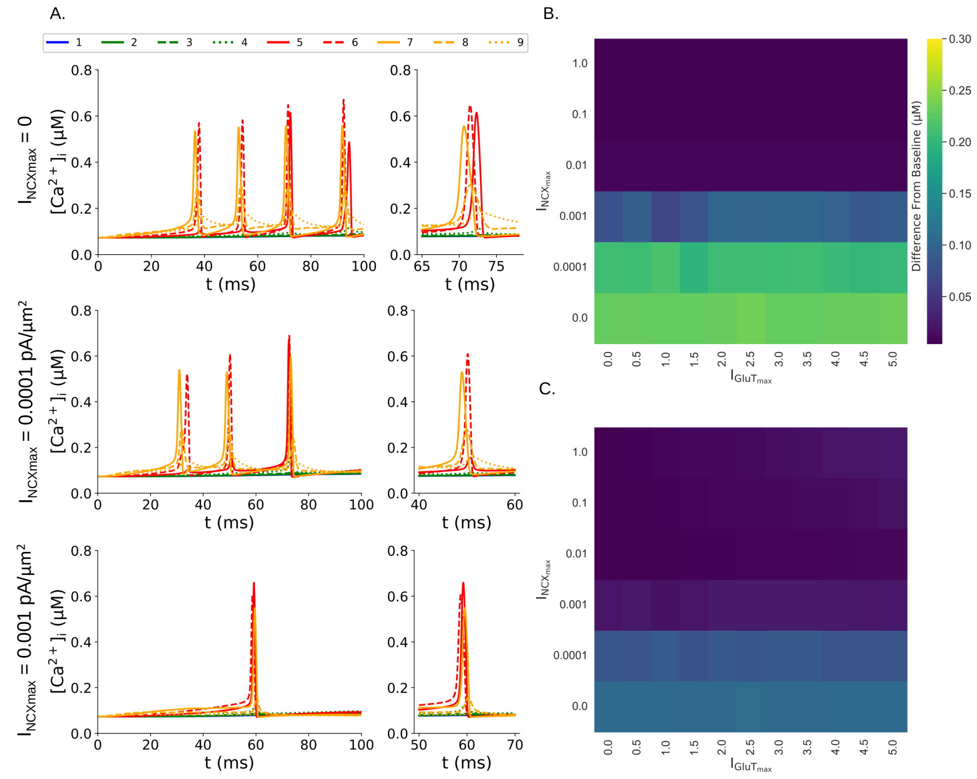


Figure 1: **Effect of GluT and NCX currents over the $Ca^{2+}$ signals in Distal Compartments.** To test the influence of the currents through GluT and NCX over the $Ca^{2+}$ dynamics, we simulated trials with combinations of different values for the parameters $J_{GluTmax}$ and $J_{NCXmax}$. In this test, compartment 9 received glutamatergic synaptic input simulated as a Poisson spike train with frequency of 10 Hz for 100 s. With $J_{NCXmax} = 0$ there were an increase in the amplitude of the $[Ca^{2+}]_i$ in distal compartments and an increase in the number of $Ca^{2+}$ signals triggered in the distal compartments when compared to the conditions with higher values of $J_{NCXmax}$. In compartment 8, NCX current is the a dominant factor regulating $[Ca^{2+}]_i$, as the variation of $J_{\mathrm{GluTmax}}$ did not change the $[Ca^{2+}]_i$ amplitude. However, in compartment 9, the increase in $J_{\mathrm{GluTmax}}$ led to a slight increase in the amplitude of $[Ca^{2+}]_i$. Since the number and amplitude of $Ca^{2+}$ are reduced by higher values of NCX current attenuates the effects of glutamatergic stimulation over the $[Ca^{2+}]_i$ and its propagation. A. $[Ca^{2+}]_i$ time series for three values of $J_{NCXmax}$ (upper: 0; middle 0.0001 pA/μm$^2$; bottom: 0.001 pA/μm$^2$). B. and C. $[Ca^{2+}]_i$ amplitudes in compartment 8 (B.) and 9 (C.)
